# Replicating the Gold Standard: A Novel Female Chronic Social Defeat Stress Model (*fem*CSDS) for Studying Sex Differences in Depression

**DOI:** 10.1101/2025.08.14.670300

**Authors:** Shashikant Patel, Roli Kushwaha, P.V Anusha, Anushka Arvind, Sainath Sunil Dhaygude, Satya Ranjan Pattnaik, Arvind Kumar, Mohammed Idris, Sumana Chakravarty

## Abstract

Depression shows significant sex differences in prevalence and neurobiological underpinnings, yet preclinical research investigating the pathophysiology of depression and the efficacy of antidepressants has predominantly relied on male models. Here, we establish a novel female chronic social defeat stress paradigm by leveraging the natural aggression of parous CD1 females, co-housed with castrated males to induce aggression while eliminating confounding sexual behaviors and without hormonal or surgical manipulations. Selected aggressive females reliably displayed offensive behaviors toward C57BL/6NCrl intruders across repeated encounters. Defeated female mice exhibited pronounced depression-like behaviors, including social withdrawal, anhedonia, behavioral despair, and elevated anxiety-like responses. Biochemical analysis revealed elevated glutamate levels in Nucleus Accumbens (NAc) and caudate putamen (CPu). Alterations in EAAT1, GRIN2B, and Neurabin expression were observed in CPu, indicating excitotoxic stress and compromised synaptic integrity. Label free Quantitative MS-MS analysis of NAc revealed 1194 significantly dysregulated proteins. Ingenuity Pathway Analysis highlighted canonical pathway disruptions in synaptogenesis signaling pathway and glutamate signaling pathway. Disease and function analysis revealed enrichment in neuroinflammation, synaptic dysfunction, and mitochondrial dysfunction. Given the extensive literature on male CSDS and its established pathophysiology, we aimed and successfully developed female-specific replica model of traditional male CSDS, enabling direct comparison and elucidation of sex differences in depression pathophysiology.

## 1. INTRODUCTION

Chronic Social Defeat Stress (CSDS), widely regarded as the gold-standard model for studying depression-like phenotypes, recapitulates the psychosocial stress, fear circuitry dysregulation, and neuroplasticity deficits observed in humans (1–3). However, its utility has been constrained by a systemic male bias. Efforts to adapt CSDS for female subjects have encountered significant obstacles rooted in sex-specific behavioral and neuroendocrine dynamics. CD1 males do not attack female intruders (4), necessitating artificial manipulations such as urine soiling on females (5,6), a method that may introduce confounding hormonal variability by altering estrous cyclicity and stress hormone profiles. Furthermore, CD1 females lack innate aggression, rendering same-sex resident-intruder paradigms unfeasible. Invasive workarounds, such as optogenetic activation of the ventromedial hypothalamus (VMHvl) in males to induce female-directed aggression, are technically fragile and lack translational relevance (7). Even in the case of experimentally manipulated male aggressors for a female CSDS setup the male-female interactions introduce confounding variables due to the olfactory cues, which can distort the experimental outcomes and reduce the validity of the findings when extrapolating or comparing data from male CSDS experiments to females.

To address these gaps, we developed a novel female CSDS (*fem*CSDS) model by strategically inducing aggression in CD1 females toward C57BL/6 females, circumventing the confounding variables inherent to male-female interactions. We hypothesized that aggression in female CD1 mice could be induced by inhibiting the evolutionary conserved reproductive instinct, analogous to retired male breeders. Unlike prior approaches that artificially manipulated male CD1 aggression towards females, our model leveraged postpartum aggression (PPA), a natural defensive behavior in lactating CD1 rodents. By co-housing parous CD1 females with castrated CD1 male partners, we prolonged PPA, inducing sustained aggression towards female C57BL/6NCrl intruders. This approach eliminates invasive manipulations and hormonal confounders, preserving ethological validity while capturing the psychosocial stress dynamics absent in traditional female depression models.

Using our paradigm, we investigated (in intruder female C57 mice) behavioral outcomes alongside neurochemical and proteomic alterations in the nucleus accumbens (NAc), a key region in the brain’s stress and reward circuitry. We demonstrate that socially defeated female mice exhibit strong depressive-like phenotypes, including social avoidance, anhedonia, and behavioral despair. Proteomic profiling of the NAc revealed extensive dysregulation in pathways governing mitochondrial metabolism, glutamate signaling, synaptic vesicle cycling, and neuroplasticity. Integrative analysis using Ingenuity Pathway Analysis (IPA) was suggestive of sex-relevant regulatory networks, pathways and proteostatic mechanisms of stress vulnerability in females.

## 2. METHODS AND MATERIAL

All animal experiments were performed in the Animal House, CSIR-CCMB, Hyderabad, India. Protocols used in this study were approved by the institutional Animal Ethics Committee under protocol numbers [#IAEC/62/2022-23] and [#IAEC 70/2022-23]. For detailed methodologies refer to supplementary methods.

### 2.1 Animals and experimental strategy

Preliminary efforts to induce aggression in intact female CD1 mice through co-housing with castrated males proved insufficient for a reliable CSDS model (refer to Supplementary for details on Experimental Workflow 1). Thus, we adopted a modified strategy by leveraging and prolonging postpartum aggression (PPA). In our modified strategy we pair housed intact male and female CD1 mice aged 3-4 months (n= 19 pairs). These pairs were co-housed for 20-30 days to allow mating, gestation, and parturition. Post parturition, the pups were sacrificed on postpartum day 2, aggravating the natural aggressive tendencies of the postpartum female. The removal of pups on day 2 allowed us to begin aggression assessments. Aggression was assessed on postpartum days 2, 3, and 4 by introducing a female C57BL/6NCrl intruder into the resident female’s (now referred to as “*parous female*”) cage. Based on the aggression assessments, we aimed at aggravating PPA, for which we extended the protocol by castrating the male CD1 partners after postpartum day 4 (Figure 1: Experimental workflow 2). Following castration, the male mice were housed individually for a week to allow for recuperation and then were reintroduced to their respective female partners. The pairs were co-housed for an additional 7-8 weeks. After this extended co-housing period, we reassessed aggression in these parous females.

**Figure 1:**
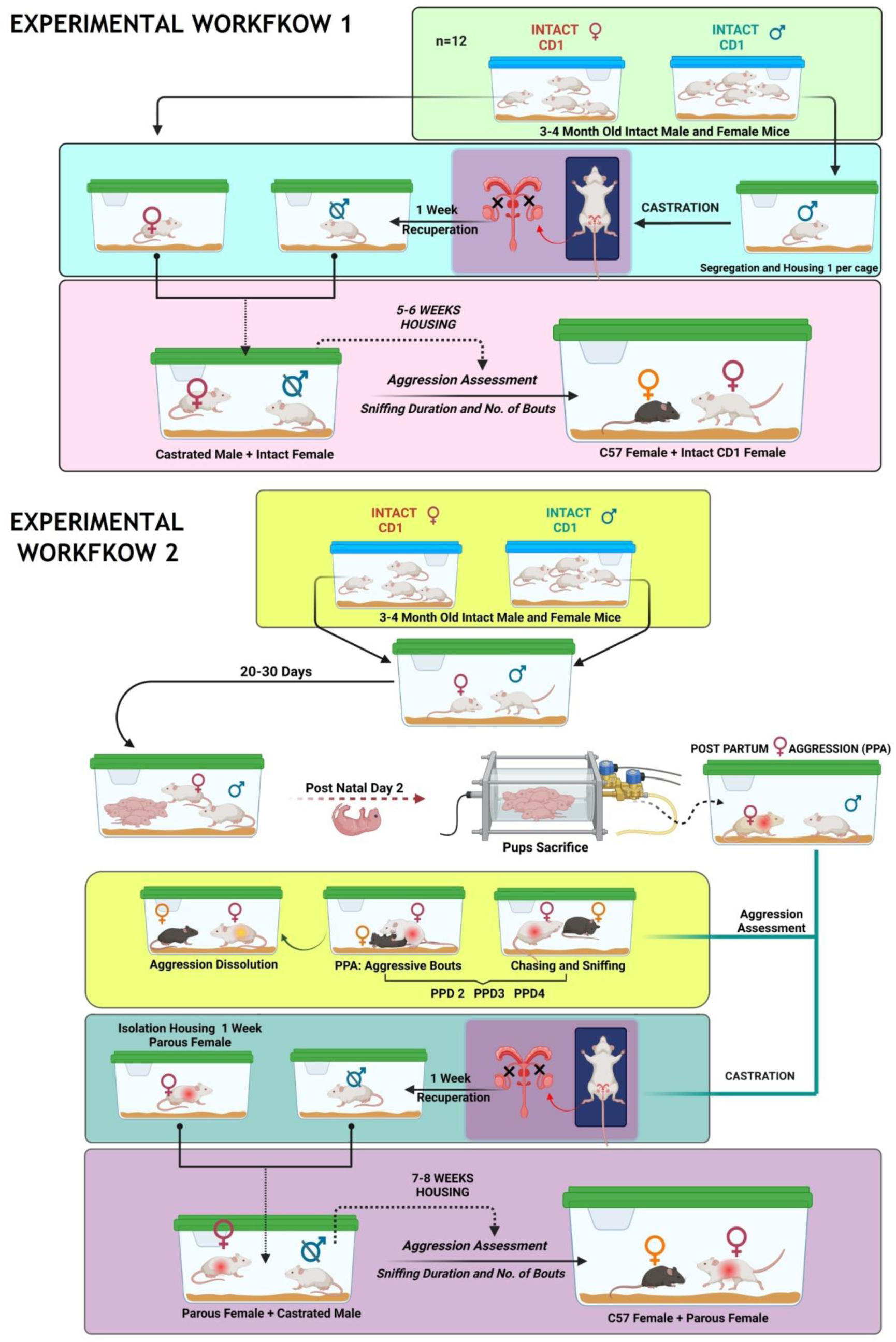
Experimental workflows for inducing and characterizing female aggression in CD1 female: This figure illustrates two distinct protocols aimed at inducing female aggression. **(A) Experimental Workflow 1: Induction of aggression in intact CD1 females;** Intact 3-4 month old CD1 male and female mice were procured and habituated. Females were then individually housed, while males were castrated and allowed to recuperate for one week. Castrated males were then co-housed with intact females for 5-6 weeks. Aggression was assessed on three alternate days by introducing a C57BL/6NCrl intruder into the CD1 female cage for 7 minutes, quantifying aggressive bouts and sniffing duration. This workflow determined if co-housing with castrated males induced aggression in intact females. **(B) Experimental Workflow 2: Enhancement and prolongation of postpartum aggression (PPA) in parous CD1 females;** Intact 3-4 month old CD1 male and female mice were pair-housed for 20-30 days to facilitate mating and parturition. On postpartum day 2, pups were sacrificed. Aggression was assessed on postpartum days 2, 3, and 4 using a C57 intruder. After assessments, the same male partners were castrated, recuperated for one week, and then co-housed with the parous female for an additional 7-8 weeks. Aggression was then reassessed on three alternate days. This workflow established sustained female aggression for *fem*CSDS studies. Selected Parous Females (SPF) were defined as those exhibiting at least one aggressive bout on all three testing days and five or more bouts on any two consecutive days.

Female aggressors for social defeat were selected based on stringent criteria; only those CD1 female mice that exhibited at least one aggressive bout on all the three testing days and achieved five or more bouts on any two consecutive days were included. For clarity and to differentiate from other female groups, these consistently aggressive mice were designated as “*Selected Parous Females” (SPF)*.

### 2.2 Female chronic social defeat stress paradigm (*fem*CSDS)

Following the selection of these SPF, a new cohort of female C57BL/6NCrl mice was procured to undergo the *fem*CSDS paradigm. These C57BL/6NCrl mice were subjected to 10 continuous days of social defeat, exposing to SPF aggressor. To avoid bias and ensure consistent stress induction, each C57BL/6NCrl mouse was shifted to a novel SPF aggressor cage. C57 females were subsequently housed overnight in the aggressor’s cage, separated by a perforated divider. Following defeat session, a behavioral battery was conducted on post-defeat day (PDD) 1, 2, and 3 to assess the development of anxiety and depression like symptoms in the intruder C57BL/6NCrl mice (Fig 3E). Post completion of paradigm the female CD1 aggressor mice were returned to their male partners for co-housing, and their aggression was reassessed approximately 4 months later, to assess long-term sustainability of their aggressive phenotype.

### 2.3 Behavioral Assessments

Aggressive behavior was evaluated in female CD1 mice as part of aggressor selection to be utilized in *fem*CSDS paradigm. Each resident-intruder interaction was recorded and analyzed by an experimenter blinded to the experimental conditions. Aggression was assessed using three parameters: sniffing duration (approach behavior), latency of first attack and number of aggressive bouts (defined as any discrete attack behavior, including chasing, biting, or pinning). The number of aggressive bouts served as the primary metric for quantifying aggression intensity. Refer supplementary file for detailed behavioral tests.

### 2.4 Preparation of tissue lysate and Label free Quantitative MS-MS analysis

Samples included 2 control groups and two defeated groups with each group consisting of pooled samples from 4 individual mice. Overall samples from 8 control and 8 defeated mice were utilized. Label free Quantitative MS-MS analysis was carried out for all the samples against control. Differential expressions in proteins were estimated relative to the non-treated control negative samples. Proteins having more than 0.5 log change were recruited for the study. Refer supplementary methods for details.

### 2.5 Statistics

All statistical analyses were conducted using GraphPad Prism (v9.0). Behavioral data were acquired and quantified using Ethovision XT 17 (Noldus, Netherlands). Data were first checked for normality using the Shapiro-Wilk test. For repeated measures data (e.g., sniffing durations or aggressive bouts across days), either repeated measures ANOVA or Friedman test was employed depending on data normality. Bonferroni or Dunn’s post hoc tests were applied where appropriate. Between-group comparisons (e.g., control vs defeated) were analyzed using unpaired two-tailed t-tests or non-parametric Mann–Whitney U tests when data did not meet parametric assumptions. One-way ANOVA or Kruskal-Wallis tests were used for comparisons among more than two groups, followed by post hoc analysis. Linear mixed-effects models (REML) were employed to account for occasional missing values. Data are presented as mean ± SEM or median with interquartile range, as appropriate. Significance was defined at p < 0.05.

## 3. RESULTS

### 3.1 Minimal aggression was observed in intact female CD1 mice

To investigate the approach behavior towards intruder C57 female, exhibited by control (CON) and intact female (IF) CD1 mice (workflow 1, Fig 1), we measured the mean sniffing duration across three alternate days (Day 1, 3 and 5). A two-way repeated measures ANOVA revealed a significant main effect of DAY (p = 0.0176), however no significant differences were observed between the groups (p = 0.5783). Bonferroni post hoc tests indicated a significant increase in sniffing duration for the IF group compared to the CON group on Day 1 (p = 0.0066). No significant differences were observed on Day 3 (p=0.2418) or Day 5 (p > 0.9999) (Fig. 2A). A Friedman test was conducted to assess differences in aggressive bout numbers between control (CON) and intact female (IF) (Fig. 2B). The test revealed no significant differences across days or between groups (p= 0.5323).

**Figure 2:**
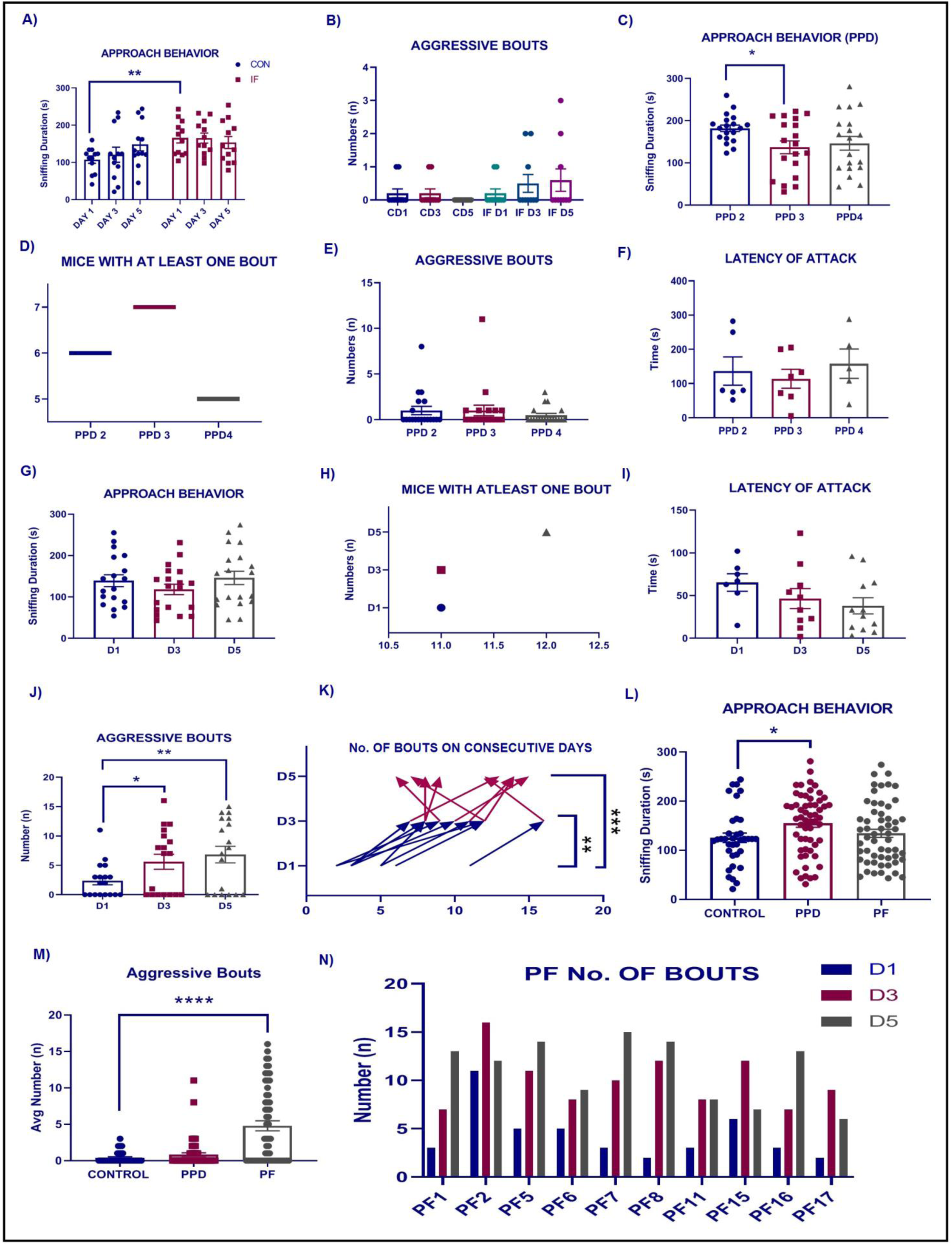
Aggression assessment in female CD1 mice across experimental workflows: **(A-B) Minimal aggression in intact CD1 females;** (A) Mean sniffing duration towards a C57BL/6 intruder in Control (CON) and Intact Female (IF) CD1 mice over Day 1, Day 3, and Day 5. A significant increase in sniffing duration was observed for the IF group compared to the CON group on Day 1. (B) Number of aggressive bouts by CON and IF groups across Day 1, Day 3, and Day 5, showing no significant differences. **(C-F) Workflow 2: Postpartum Aggression (PPA) Alone;** (C) Mean sniffing duration in parous CD1 females on postpartum days 2 (PPD 2), 3 (PPD 3), and 4 (PPD 4). A significant decrease in sniffing duration was observed from PPD 2 to PPD 3. (D) Number of parous mice exhibiting at least one aggressive bout on PPD 2, PPD 3, and PPD 4. (E) Total number of aggressive bouts by parous CD1 females on PPD 2, PPD 3, and PPD 4, indicating no significant changes across days. (F) Latency of attack in parous CD1 females on PPD 2, PPD 3, and PPD 4, showing no significant differences. **(G-J) Workflow 2: Aggression Post Co-habitation with Castrated Male Partner;** (G) Mean sniffing duration in PF towards a C57BL/6 intruder on Day 1 (D1), Day 3 (D3), and Day 5 (D5) after extended co-housing, showing consistency across days. (H) Number of PF mice exhibiting at least one aggressive bout on D1, D3, and D5. (I) Latency of attack in PF mice on D1, D3, and D5. One-way ANOVA indicated no significant differences in latency, although a trend for reduced latency was observed. (J) Total number of aggressive bouts by PF mice on D1, D3, and D5, showing a significant increase from D1 to D3 and D1 to D5. **(K-N) Comparative analysis of SPF;** (K) Number of aggressive bouts exhibited by SPF over D1, D3, and D5, demonstrating a significant increment in aggressiveness. (L) Cumulative sniffing duration comparison across Control, PPD, and PF groups. PPD mice showed significantly longer sniffing duration than controls. (M) Cumulative number of aggressive bouts across Control, PPD, and PF groups. PF mice exhibited a significantly higher number of bouts compared to controls. (N) Individual aggressive bout counts for SPF (out of 19 PF) across D1, D3, and D5. Data are shown as mean ± SEM or individual data points. Statistical significance is indicated as *p < 0.05, **p < 0.01, ***p < 0.001, ****p < 0.0001.

### 3.2 Postpartum aggression alone is insignificant for successful implementation in CSDS modeling

In the modified protocol designed to assess PPA, we conducted aggression assessments at two stages. The first assessment was performed after parturition on postpartum days 2 (PPD 2), 3 (PPD 3), and 4 (PPD 4). Repeated measures ANOVA was conducted where Bonferroni post hoc comparisons showed a significant decrease in sniffing duration from PPD 2 to PPD 3 (p = 0.0228). No significant difference was observed between PPD 2 and PPD 4 (p = 0.1145) (Fig 2C). Importantly, of the 19 mice evaluated, only 6 exhibited at least one bout on PPD 2, while 7 did so on PPD 3 and only 5 on PPD 4 (Fig 2D). Furthermore, only 3 mice demonstrated at least one bout across all three days of assessment. The Friedman test indicated no significant changes in bout no’s across these days (p = 0.3813) (Fig 2E). It is noteworthy that the latencies of aggression could only be plotted for mice which exhibited bouts during the observations. Kruskal-Wallis test showed no significant differences in attack latencies between postpartum days (P = 0.6695) (Fig 2F).

### 3.3 Aggression assessment in parous females post co-habitation with castrated male partner

We reassessed the aggression to investigate whether prolonged postpartum habituation with castrated male partners could enhance aggressive behavior. The PF were tested for aggression towards C57 mice on three alternate days: Day 1, 3 and 5. A linear mixed-effects model (REML) was applied to account for one missing value on D1 (Duration of sniffing and number of bouts of one animal out of 19 couldn’t be recorded due to experimental error). Bonferroni’s multiple comparisons confirmed that neither D1 vs. D3 (p = 0.7552) nor D1 vs. D5 (p > 0.9999) differed significantly. These findings indicate that approach behavior remained consistent across the testing days (Fig 2G).

Among the 19 female mice assessed, 11 displayed at least one bout on D1 and D3, while 12 mice exhibited on D5. 10 mice displayed on all the three days of testing (Fig 2H). Latency to attack in these mice was assessed using a one-way ANOVA. The analysis indicated no significant differences in latency to attack between the three days (F(2, 26) = 1.507, p = 0.2402). Post-hoc Bonferroni comparisons revealed no significant differences between D1 and D3 (p = 0.5205) or between D1 and D5 (p = 0.1902) (Fig 2I). Friedman test revealed a significant effect of the testing day on the number of aggressive bouts (χ²(2) = 18.57, p < 0.0001). Dunn’s multiple comparisons test indicated a significant increase in aggressive bouts from D1 to D3 (p = 0.0313) and further from D1 to D5 (p = 0.0017). Note that we calculated the total number of aggressive bouts across all assessed days, irrespective of the differential number of individual mice that gave bouts on specific days of testing (Fig 2J).

### 3.4 Aggression assessments in selected parous female CD1 mice

The Friedman test revealed a highly significant difference among the days (Friedman statistic = 15.85, p < 0.0001). Dunn’s multiple comparisons test further indicated a significant increase in aggressive bouts on D3 compared to D1 (rank sum difference = -13.50, adjusted p = 0.0051) and on D5 compared to D1 (rank sum difference = -16.50, adjusted p = 0.0004) (Fig 2K). These findings demonstrate a clear, incrementing trend in aggressiveness over the testing period.

### 3.5 Comparative Analysis of Aggressive Behavior in Control, PPD, and PF Groups

In comparing cumulative sniffing duration across all the three testing days in control, PPD, and PF mice, a Kruskal-Wallis test indicated a significant difference among the groups (Kruskal-Wallis statistic = 6.185, approximate p = 0.0454). Dunn’s post-hoc test revealed that PPD mice had a significantly longer sniffing duration than controls (adjusted p = 0.0487), while no significant difference was found between PPD and PF groups (adjusted p > 0.9999) (Fig 2L). Kruskal-Wallis test revealed a highly significant overall difference in the number of bouts among the groups (Kruskal-Wallis statistic = 26.15, p < 0.0001). Post-hoc Dunn’s multiple comparisons test showed a significant increase in the number of bouts in PF mice compared to controls (adjusted p < 0.0001), while no significant difference was observed between control and PPD groups (adjusted p = 0.9897) (Fig 2M). The number of aggressive bouts by individual selected parous females on testing days is presented in Fig 2N.

### 3.6 Comparative Analysis of Aggressive Behavior in Control and SPF Groups

We compared SPF group with normal CD1 females (Control) to decipher the aggression levels in selected aggressors compared to un-manipulated female CD1 mice. Mann–Whitney U test was performed on sniffing duration data indicating no significant difference in approach behavior between the groups (Fig 3A). In contrast, comparison of cumulative aggressive bout numbers over three testing days showed a marked difference between the groups. A Mann–Whitney U test yielded U = 7 with an exact p-value < 0.0001, demonstrating a highly significant increase in aggressive bouts in the SPF group relative to controls (Fig 3B).

**Figure 3:**
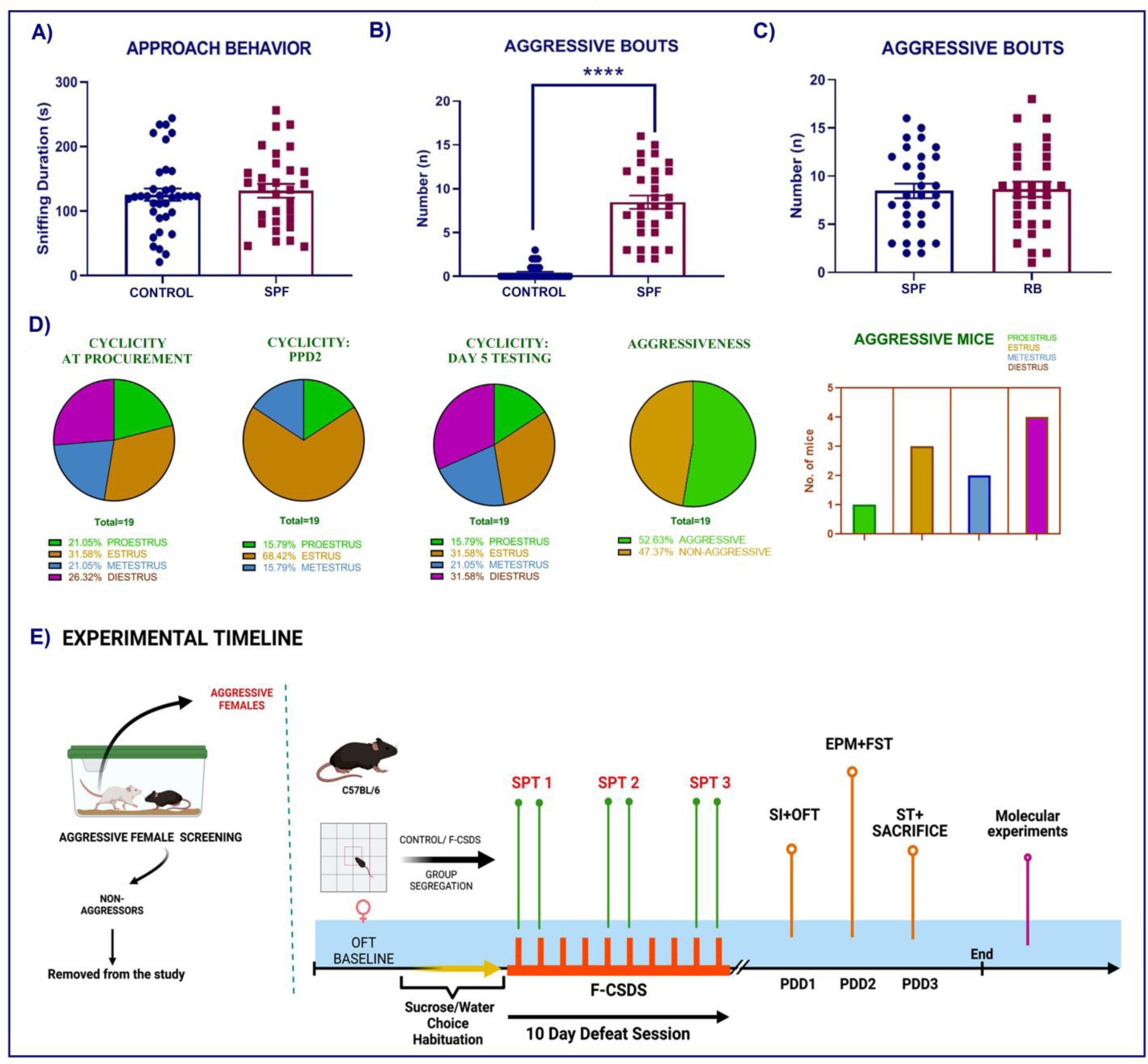
Comparative analysis of aggressive behavior in SPF, cyclicity in female CD1 mice and *fem*CSDS timeline: This figure presents a comprehensive comparative analysis of aggression levels in selected parous females (SPF) relative to control females and male retired breeders, along with the influence of estrous cyclicity on aggressive behavior. **(A-B) Comparative Aggression: Control vs. SPF;** (A) Sniffing duration (s) towards a C57BL/6 intruder in Control vs. SPF CD1 females. No significant difference was found in approach behavior (p = 0.5623). (B) Cumulative aggressive bouts over three days for Control vs. SPF groups. SPF mice showed significantly more aggressive bouts than controls (p < 0.0001), indicating robust aggression. (C) Aggression Levels: SPF vs. Male Retired Breeders. No significant difference was observed (p = 0.8802), suggesting SPF aggression is comparable to established male aggressors. (D) Estrous Cyclicity and Aggression: Pie charts show estrous cyclicity distribution in female CD1 mice at procurement (21.05% Proestrus, 31.58% Estrus, 21.05% Metestrus, 26.32% Diestrus; n=19), on Postpartum Day 2 (PPD 2) (15.79% Proestrus, 68.42% Estrus, 15.79% Metestrus; n=19), and on Day 5 of testing (15.79% Proestrus, 31.58% Estrus, 21.05% Metestrus, 31.58% Diestrus; n=19). The bar graph shows 52.63% of parous females were aggressive (SPF), with most aggressive mice (4 of 10) in diestrus, others distributed across proestrus (1), estrus (3), and metestrus (2). (E) Experimental Timeline: After screening for aggressive females, selected CD1 mice (SPF) were used to induce chronic social defeat in a new set of C57BL/6 female mice. C57BL/6 mice underwent baseline assessments and habituation to a sucrose/water two-bottle choice setup for the Sucrose Preference Test (SPT). The *fem*CSDS paradigm involved 10 continuous days of social defeat sessions. During this 10-day period, sucrose preference was measured on Days 1, 2, 5, 6, 9, and 10 to monitor anhedonic behavior. Following the 10-day defeat period, a behavioral battery was conducted on Post Defeat Days 1, 2, and 3 to assess anxiety and depression like behaviors. This battery of behavioral tests included SIT, OFT, EPM, FST and SST. Sacrifice was done on PDD 3 after the last behavioral test. ****p < 0.0001, n=10 SPF.

### 3.7 Aggressive Behavior in SPF is comparable to that of CD1 male retired breeders

To explore the feasibility of using SPF as aggressors, we compared them with a separate set of retired male breeders. The t-test showed no significant difference in the mean bout counts between the two groups (t = 0.1514, df = 58, p = 0.8802). F-test comparing variances also found no significant difference (F(29, 29) = 1.103, p = 0.7942), confirming homogeneity of variance across groups. These findings demonstrate that SPF mice display an aggressive phenotype that is statistically similar to that of traditional male retired breeders (Fig 3C).

### 3.8 Effect of cyclicity on aggressive behavior in mice

Significant aggression was induced in 10 out of 19 females (52.63%). Analysis of estrous cyclicity revealed variations in the proportion of females at different stages across the experimental timeline. At procurement, the distribution was relatively balanced, with 21.05% in proestrus, 31.58% in estrus, 21.05% in metestrus, and 26.32% in diestrus. Notably, the aggressive SPF mice were predominantly in the diestrus stage (4 out of 10), indicating a potential correlation between this phase and increased aggression (Fig 3D).

### 3.9 Behavioral Assessment of Female C57 Mice Subjected to Chronic Social Stress

In SIT, defeated mice displayed significantly lower levels of social interaction compared to control mice (p = 0.0045) (Fig 4A). In evaluating the frequency of interaction zone entries, no significant difference was found between defeated and control mice (Fig 4B). In OFT the defeated group exhibited significantly lower median time in the central zone (1.511%) compared to the control group (4.611%), indicating an increase in anxiety-like behavior (p = 0.0106) (Fig 4D). Unpaired t-test indicated no significant difference in the distance traversed between the two groups (p = 0.1623) (Fig 4E) while significant reduction in the average velocity was observed (Fig 4F) alongside reduction in the central zone frequency in the defeated group (p = 0.0391) (Fig 4H).

**Figure 4:**
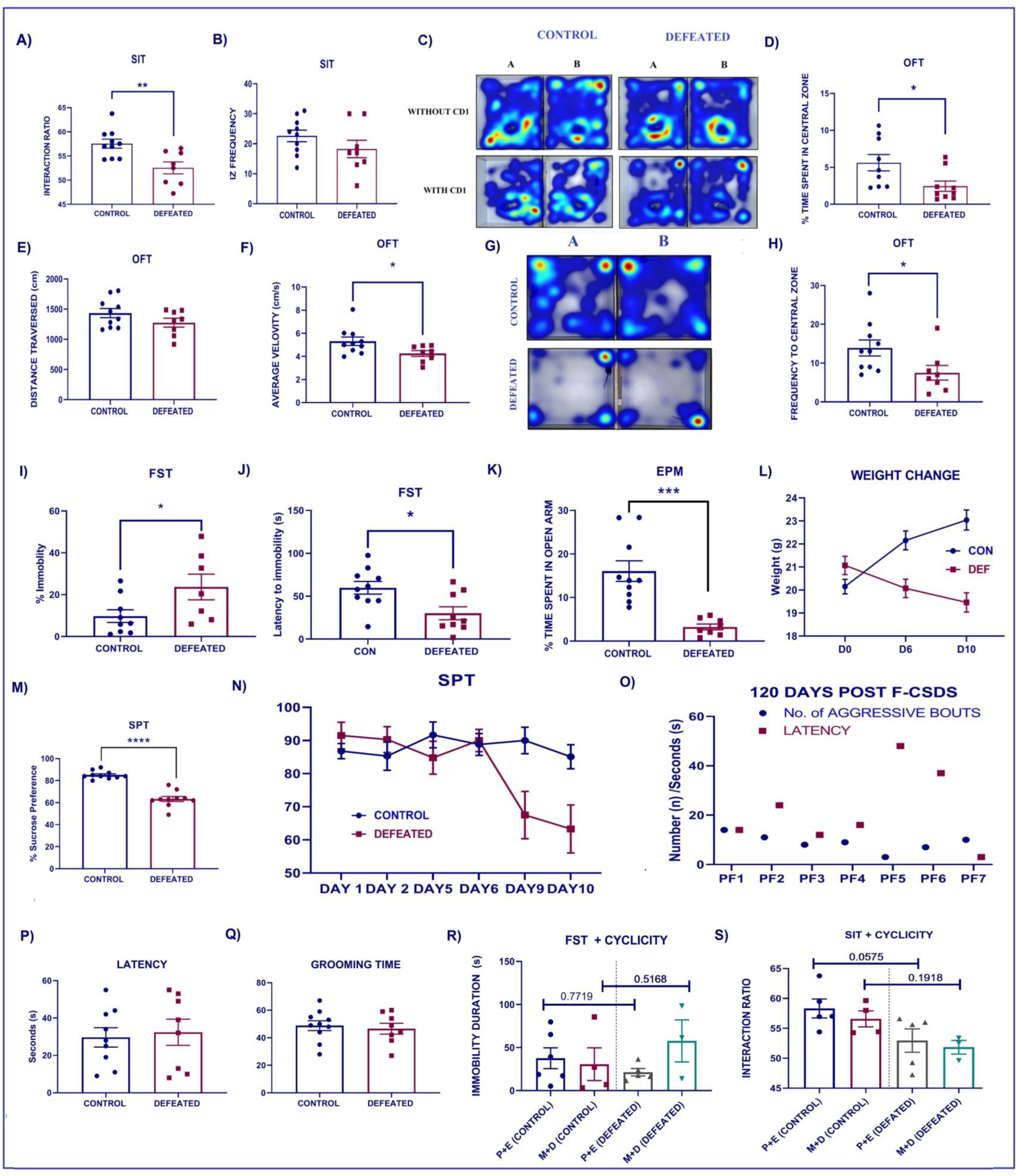
Behavioral and physiological consequences of *fem*CSDS in C57BL/6 mice. In SIT, defeated mice exhibited significantly lower interaction ratios (A) compared to controls, indicating social avoidance, though interaction zone entry frequency was unchanged (B); accompanying heat maps illustrate representative exploration patterns during the SIT without and with a CD1 aggressor (C). OFT indicated induction of anxiety like behavior in defeated mice, evidenced by significantly reduced time spent in the central zone (D). While total distance traversed (E) showed no significant difference, average velocity was reduced (F). Representative heatmap of OFT show activity distribution (G). Furthermore, defeated mice displayed lower central zone frequency (H). In the FST, defeated mice displayed hallmarks of behavioral despair, showing significantly increased immobility time (I) and reduced latency to immobility (J). Anxiety-like behavior was further visualized in EPM, where defeated mice spent significantly less time in the open arms (K). Physiologically, defeated mice experienced significant body weight loss by Day 10 of defeat compared to controls (L). Anhedonia, a key depressive symptom, was robustly evident in defeated mice, demonstrating significantly lower sucrose preference on Day 10 of the SPT (M), with a progressive decline observed across all assessment days (N). The aggressive phenotype of the CD1 female aggressors proved highly stable, with reassessments 120 days post *fem*CSDS showing that most maintained robust aggressive bouts comparable to initial levels, suggesting their reusability (O). In the SST, no significant differences were observed between defeated and control mice in latency to initiate grooming (P) or total grooming duration (Q). Finally, when evaluating the influence of estrous cyclicity, neither FST immobility duration (R) nor SIT interaction ratios (S) showed significant modulation by cycle stage (Proestrus + Estrus vs. Metestrus + Diestrus) in either control or defeated mice, though a trend towards reduced interaction was noted in P+E defeated vs. P+E control groups. All data are presented as mean ± SEM or individual data points with statistical significance indicated as *p < 0.05, **p < 0.01, ***p < 0.001, ****p < 0.0001.

Defeated mice exhibited a significant increase in FST immobility time compared to controls, indicative of behavioral despair (p = 0.0468) (Fig 4I). The latency to immobility was significantly reduced (p = 0.0122) (Fig 4J). Defeated mice exhibited reduced exploration of open arms compared to controls in EPM (p = 0.0002) (Fig 4K). Bonferroni’s post hoc analysis showed significant weight loss in the defeated group compared to controls at D6 (p = 0.0060) and D10 (P < 0.0001) (Fig 4L). Defeated mice at day 10 exhibited a significantly lower sucrose preference (p < 0.0001) (Fig 4M). The SPT measurements for all assessed days as plotted in Fig 4N indicated progressive decline in the preference to sucrose in the defeated mice.

After completion of the CSDS paradigm the female CD1 aggressor mice were co-housed again with their male partners. We reassessed their aggression towards C57BL/6 mice after around 120 days (4 months) over 3 consecutive testing days (Day 1, Day 2, and Day 3). The results plotted for Day 3 of testing showed that out of the 9 aggressors tested 7 of those exhibited a significant number of aggressive bouts, comparable to their initial level of aggression suggesting that aggression was still robust (Fig 4O).

In the sucrose splash test, no significant differences were observed between controls and defeated female mice in either the latency to initiate grooming (Fig 4P) or the total grooming duration (Fig 4Q). To determine whether the estrous cycle stage modulates behavioral outcomes, we stratified control and defeated C57BL/6 females based on their cycle stage, proestrus + estrus (P+E) or metestrus + diestrus (M+D) and compared in FST and SIT. In the FST, immobility duration did not significantly differ across cycle stages in either control or defeated mice (P+E vs M+D in controls: p = 0.7719; P+E vs M+D in defeated: p = 0.5168) (Fig 4R). Similarly, in the SIT, interaction ratios showed no significant differences across cycle stages in defeated mice in the M+D stage (p = 0.1918) and P+E (p = 0.0575) (Fig 4S).

### 3.10 Glutamate excitotoxicity signatures in the Caudate Putamen and Nucleus Accumbens

Immunoblotting analysis revealed a robust upregulation of NMDAR2B expression in the caudate putamen (CPu) of defeated females compared to controls (p < 0.0001, Fig. 5A), suggesting enhanced glutamatergic excitability. In contrast, the expression of the astrocytic glutamate transporter EAAT1 was significantly reduced (p < 0.01, Fig. 5B), implicating compromised glutamate clearance. Additionally, Neurabin, a postsynaptic scaffolding protein, was markedly downregulated in defeated animals (p < 0.0001, Fig. 5C), suggesting potential synaptic instability (Fig. 5H). Quantification using glutamate ELISA kit, we observed significantly elevated glutamate levels in the CPu (p < 0.01, Fig. 5D) and NAc (p < 0.05, Fig. 5E) of defeated females, aligning with the observed molecular changes. Serum glutamate levels did not differ significantly between groups (Fig. 5F), suggesting that the observed alterations were brain region-specific and not reflective of systemic glutamate changes. Furthermore, consistent with stress-associated endocrine activation, serum corticosterone levels were significantly elevated in defeated females (p < 0.01, Fig. 5G).

**Figure 5:**
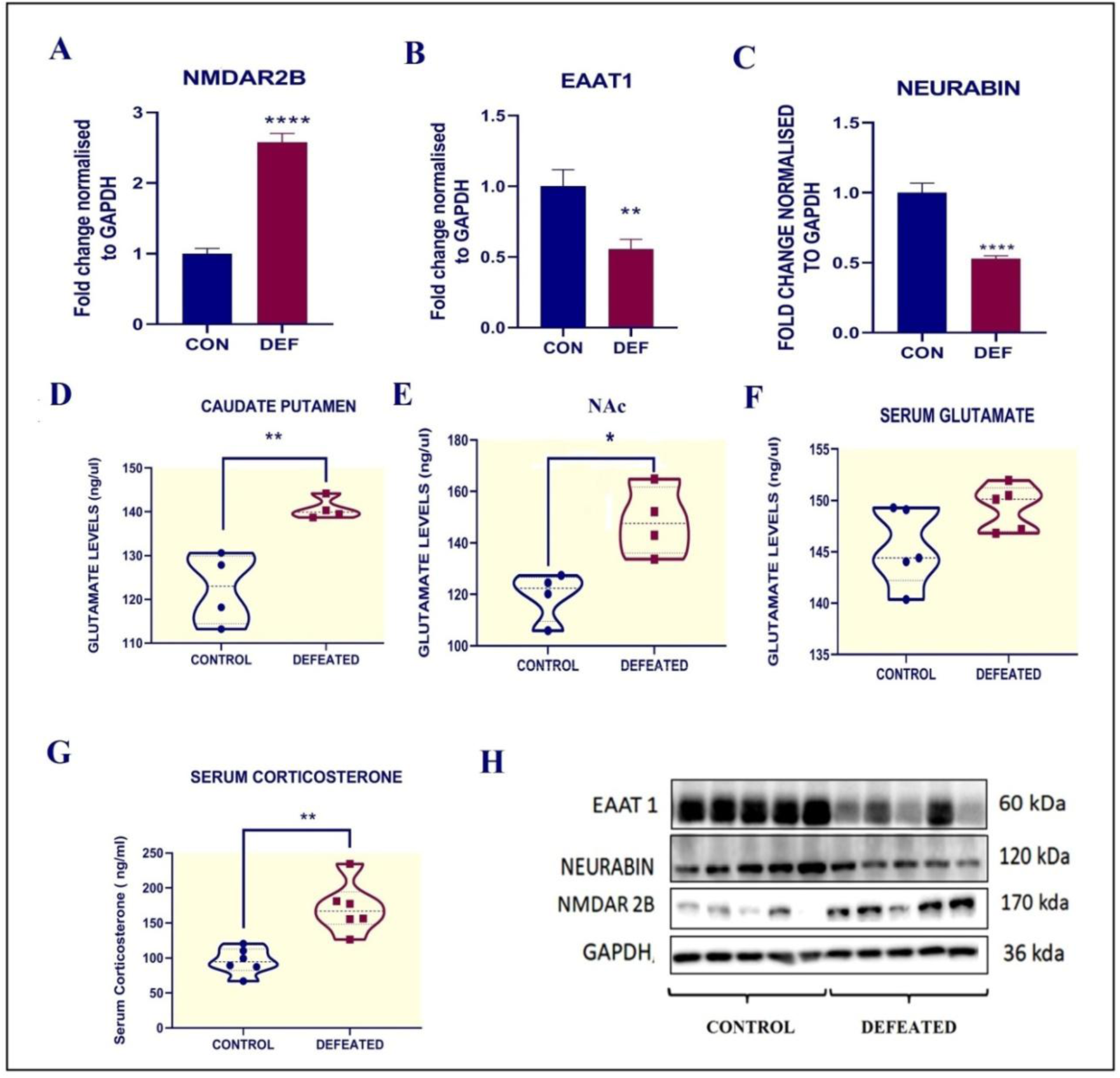
Glutamatergic Dysregulation and HPA Axis Activation in Female C57BL/6 Mice Following Chronic Social Defeat Stress. Western blot analysis of the CPu revealed a robust upregulation of NMDAR2B (A), suggesting enhanced neuronal excitability, alongside a significant reduction in the astrocytic glutamate transporter EAAT1 (B), implicating compromised glutamate clearance from the synaptic cleft. Further, Neurabin, a key postsynaptic scaffolding protein, was markedly downregulated (C), indicative of synaptic instability; these molecular changes are visually corroborated by representative immunoblots (Fig. 5H). Correspondingly, glutamate levels were significantly elevated in both the CPu (D) and NAc (E), directly confirming regional excitotoxicity, while serum glutamate levels remained unchanged (F). Consistent with chronic stress, serum corticosterone levels were significantly elevated in defeated females (G), indicating robust HPA axis activation.

### 3.11 MS/MS Proteomic landscape of the Nucleus accumbens in defeated female mice

To elucidate the molecular underpinnings of CSDS in female mice, we performed Ingenuity Pathway Analysis (IPA) on proteomic data from the NAc. A total of 3004 proteins were identified (Refer Supplementary excel). Among these, 521 proteins (17.3%) were significantly upregulated, while 673 proteins (22.4%) were downregulated in the defeated group compared to controls. A summary of IPA analysis is presented in figure 6.

**Figure 6.**
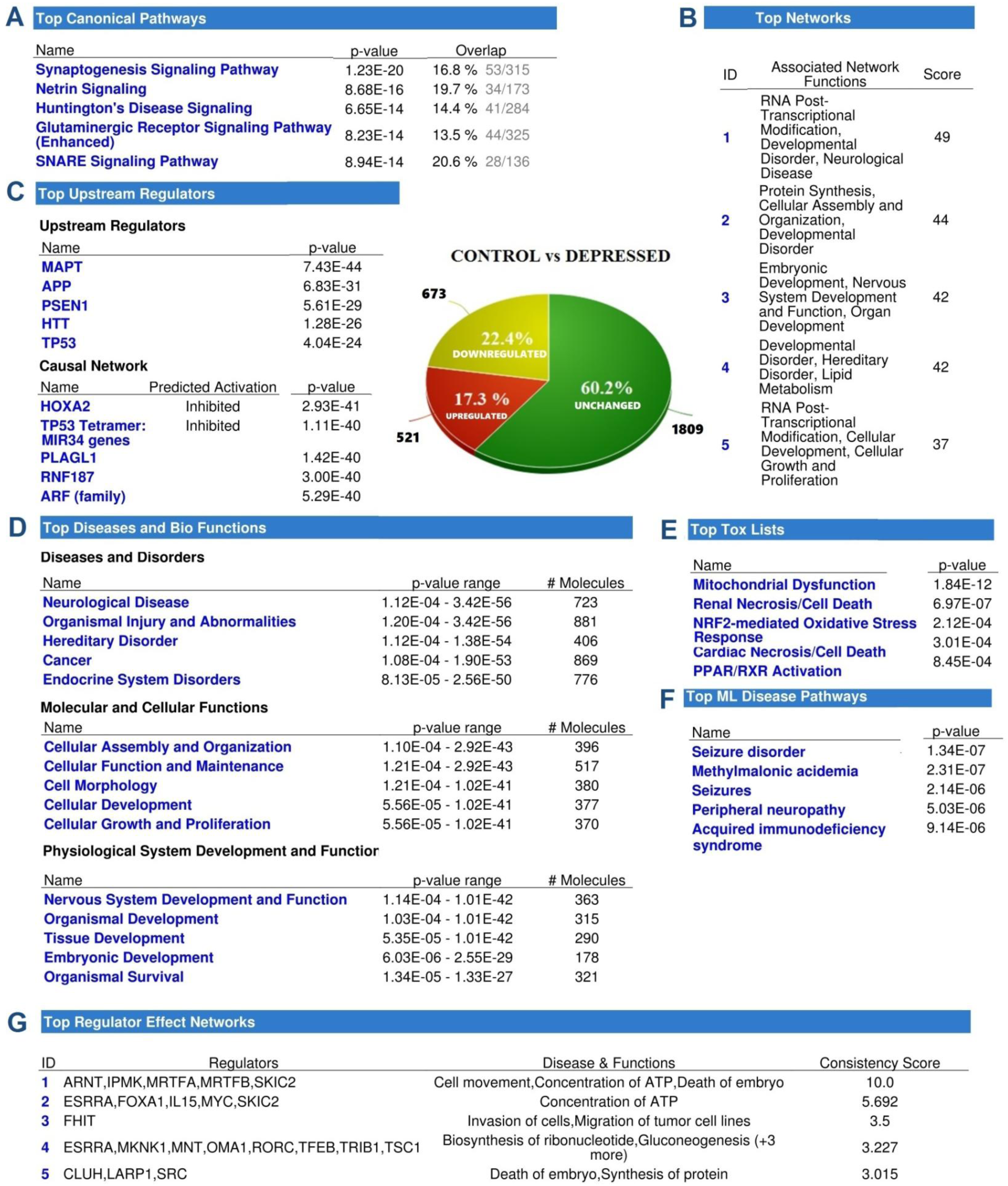
Summary of the IPA results comparing expression profiles between control and depression groups. A) Top Canonical Pathways highlights significantly enriched canonical pathways, including Synaptic Neurotransmission, Netrin Signaling, and Huntington’s Disease Signaling. (B) Top Networks displays the top five most significant molecular networks, ranked by score, and details their associated functions. (C) Top Upstream Regulators lists the most significant predicted upstream regulators and components of the causal network, along with their p-values and predicted activation states. (C) Represents the proportion of significantly downregulated (22.4%), unchanged (17.3%), and upregulated (60.2%) proteins based on proteomics data. (D) Top Diseases and Bio Functions presents enriched categories for Diseases and Disorders, Molecular and Cellular Functions and Physiological System Development and Function. (E) Top Tox Lists shows significant toxicology-related functions such as Mitochondrial Dysfunction. (F) Top ML Disease Pathways indicates relevant disease pathways from literature, including Seizure disorder. (G) Top Regulator Effect Networks identifies influential regulator networks (ARNT, JPMK, MRTFA, MRTFB, SKIC2) and their associated diseases.

#### 3.11.1 Canonical Pathway Analysis: Synaptic, Metabolic, and Neurodegenerative Dysregulation in *fem*CSDS

IPA analysis suggested extensive perturbations across several canonical pathways, spanning synaptic plasticity, neurodegeneration, metabolic homeostasis, and inflammatory signaling. The most significantly enriched canonical pathways were associated with synaptic plasticity and neurodegeneration (Fig 6A). The Synaptogenesis Signaling Pathway (*p* = 1.23E-20, 16.8% overlap) emerged as the top altered pathway (Table Sup_1). List of some of the major canonical pathways altered is presented in figure 7A. Figure 8 represents the summarized perturbations in canonical pathways (Fig 8A) along with enriched Disease and Biofunctions (Fig 8 B, C). Enrichment of the glutamatergic receptor signaling pathway (Fig 9) was marked by significant molecular alterations. Notably, downregulation was observed for the glutamate transporter SLC1A3, the metabotropic glutamate receptor GRM2 and several G protein subunits including GNG7, GNG12 and GNB5 along with glutaminase (GLS). These changes collectively suggest compromised astrocytic glutamate clearance and dysregulation in GPCR mediated glutamatergic signaling. Conversely, AMPA receptor subunits GRIA2 and GRIA4 were upregulated, potentially indicating altered postsynaptic excitability or compensatory mechanisms (Table 1). This overall pattern of dysregulation in glutamate transport and receptor signaling likely contributes to extracellular glutamate accumulation and excitotoxic damage.

**Figure 7.**
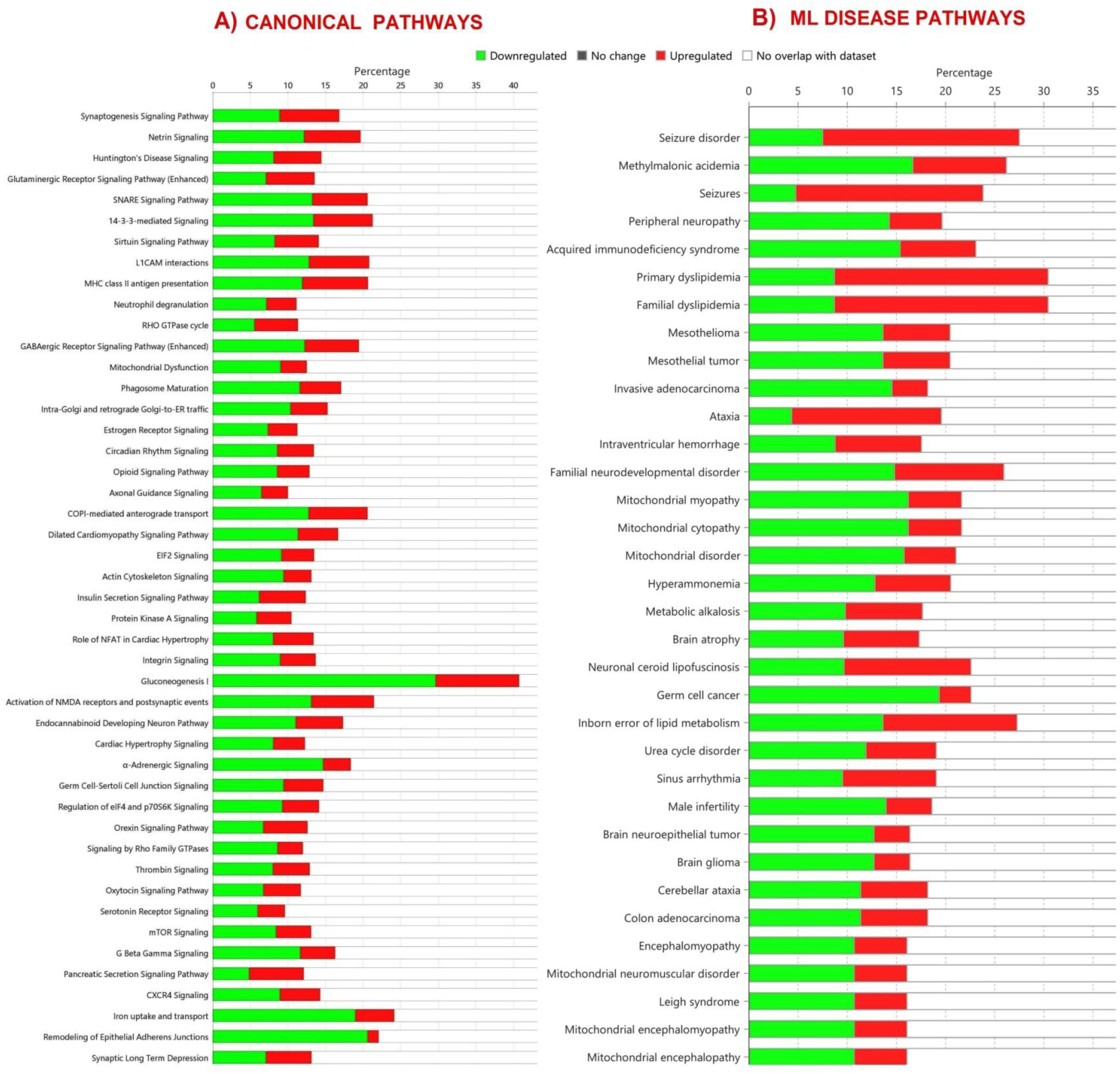
IPA analysis of canonical pathways and ML disease pathways. (A) Canonical Pathways displays a list of some of the major altered canonical pathways, indicating extensive perturbations across synaptic plasticity, neurodegeneration, metabolic homeostasis, and inflammatory signaling. (B) Top ML Disease Pathways presents the most relevant ML (Machine Learning) disease pathways, many of which are associated with neurodegenerative and neurodevelopmental disorders, indicating an acceleration of neurodegenerative processes by chronic stress.

**Figure 8:**
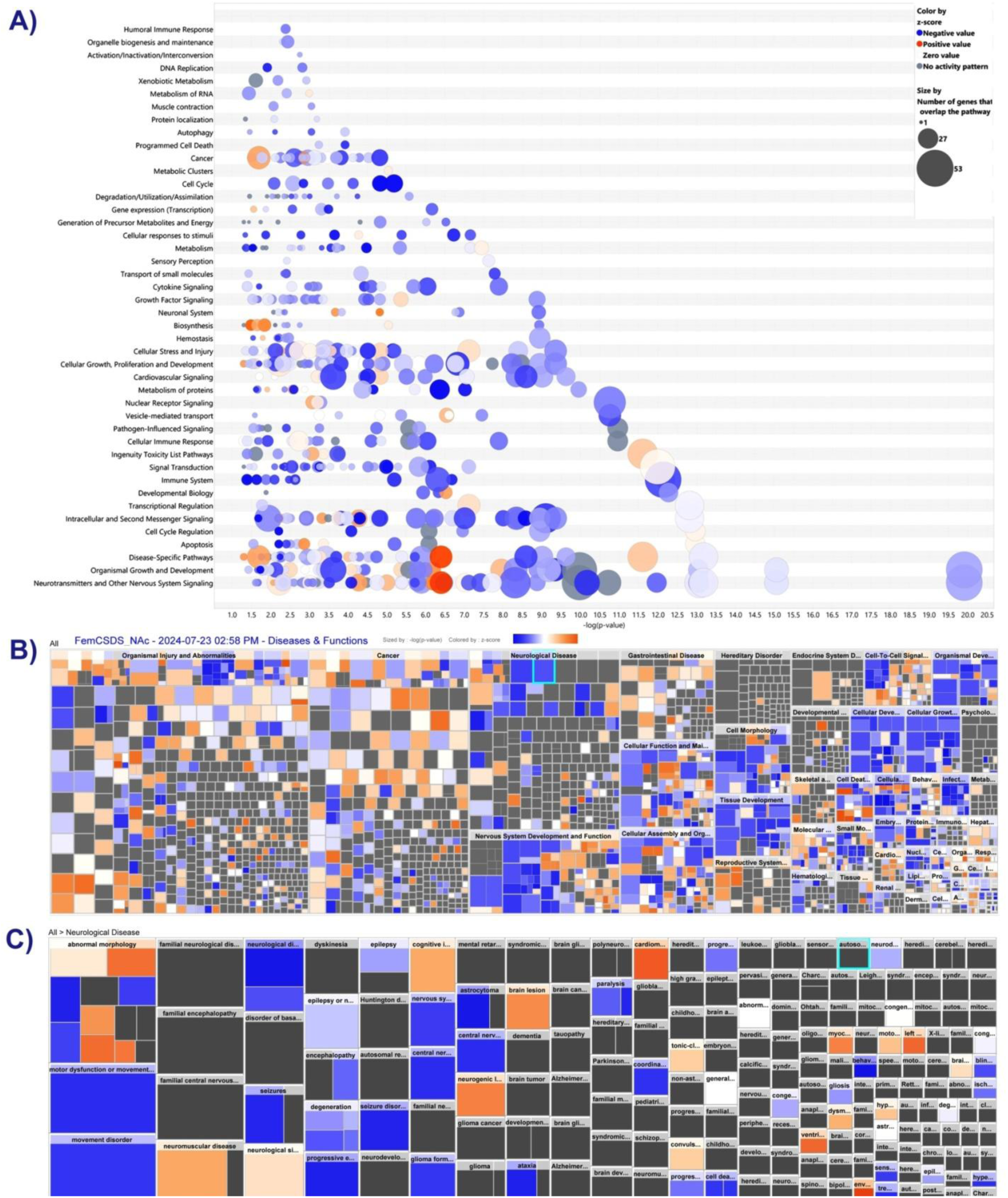
Canonical pathway summary representative bubble graph and Diseases and Biofunctions Heatmaps. **(A)** The bubble graph hierarchically visualizes canonical pathway dysregulation. Each bubble corresponds to a pathway, with its size often correlating with the number of molecules from the dataset in that pathway, and its color or position indicating the predicted activation state (activated or inhibited) or significance. Key perturbed pathways include Synaptogenesis Signaling Glutamatergic Receptor Signaling, Mitochondrial Dysfunction, and others like SNARE Signaling and TCA Cycle. **(B) Heatmap of Top Diseases and Biofunctions (All Categories)**: Comprehensive heatmap of the most significantly altered disease and biofunction categories across various cellular and physiological processes. The intensity of the color in each cell reflects the level of enrichment.**(C) Heatmap of Neurological Disease and Biofunctions**: This panel provides a focused heatmap specifically on neurological diseases and functions. It highlights the most impacted neurological categories, allowing for a detailed view of the neurobiological consequences of chronic stress.

**Figure 9.**
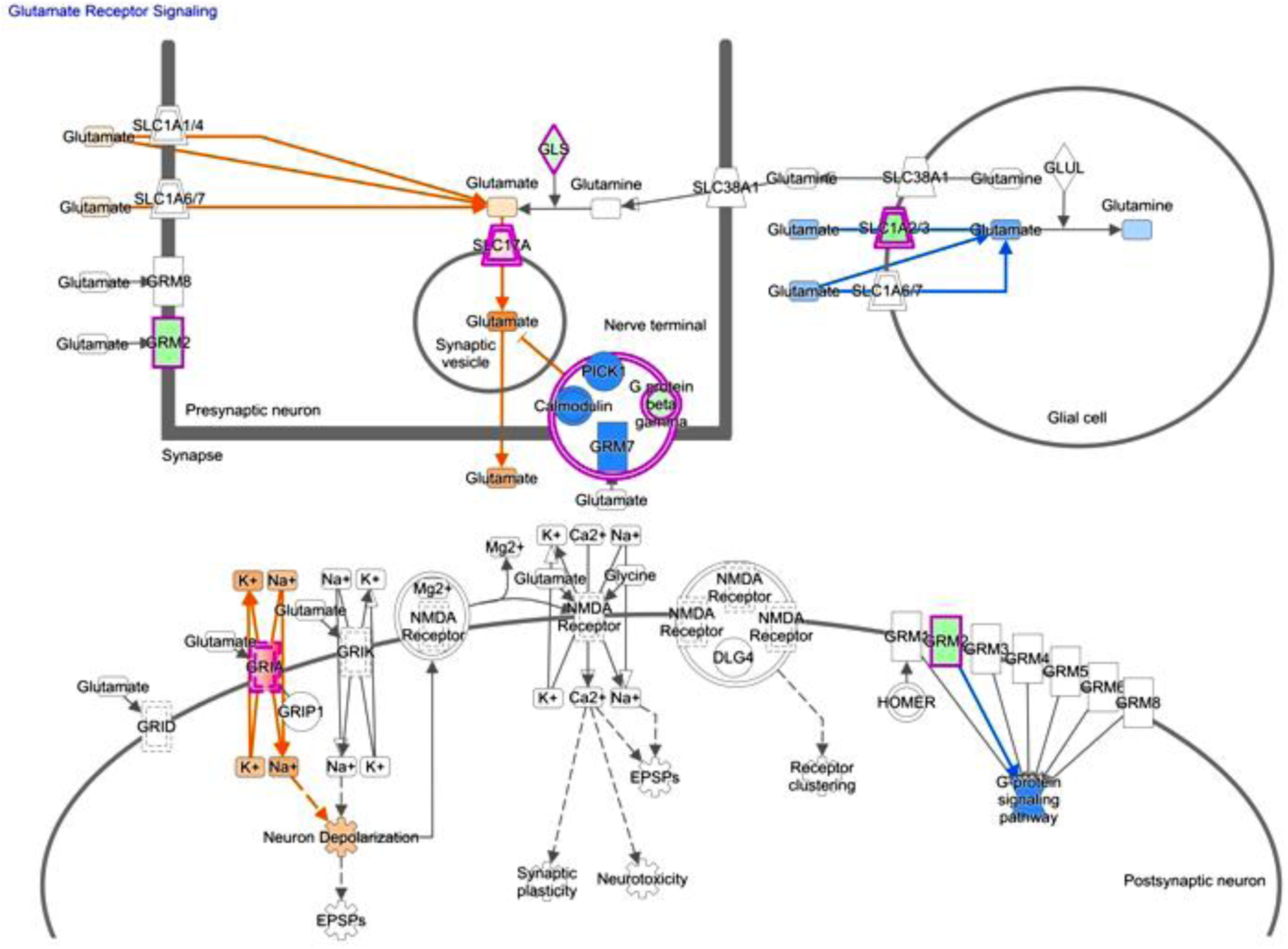
Glutamatergic Receptor Signaling Pathway. The pathway map illustrates the network of proteins governing excitatory neurotransmission, with molecules colored red indicating upregulation and green indicating downregulation in depressed females. Uncolored molecules were detected but not significantly altered. Interactions are shown with solid lines for direct and dashed lines for indirect relationships, while orange arrows predict activation and blue arrows predict inhibition of downstream effects. The analysis reveals key dysregulations in this pathway. Notably, the glutamate transporter SLC1A2 (green) is downregulated, suggesting impaired glutamate reuptake and potential extracellular accumulation. Concurrently, the GRIN2A (green) subunit of the NMDA receptor is reduced, indicating compromised NMDA receptor-mediated signaling vital for synaptic plasticity. While GRIA2 (red) and GRIA4 (red) (AMPA receptor subunits) show upregulation, potentially as a compensatory mechanism, other crucial components like GRM2 (green) and various G-protein subunits (GNG7, GNG12, GNB5 - all green) are downregulated. These collective alterations indicate a profound imbalance in the glutamatergic system, contributing to altered synaptic function, reduced plasticity, and compromised neuronal excitability.

**Table 1:** Key molecules in the Glutamatergic Receptor Signaling Pathway with their expression log ratios identified through IPA analysis.

| Symbol | Entrez Gene Name | GenPept/RefSeq | Expr Log Ratio | Expected | Location | Type(s) |
| --- | --- | --- | --- | --- | --- | --- |
| SLC1A3 | solute carrier family 1 member 3 | AAK01708.1 | -2.175 |  | Plasma Membrane | transporter |
| GRM2 | glutamate metabotropic receptor 2 | EDL21160.1 | -1.888 | Up | Plasma Membrane | G-protein coupled receptor |
| GNG7 | G protein subunit gamma 7 | XP_036011512.1 | -1.424 |  | Plasma Membrane | other |
| GNG12 | G protein subunit gamma 12 | CAG9553592.1 | -1.272 |  | Plasma Membrane | enzyme |
| GNB5 | G protein subunit beta 5 | NP_034443.1 | -1.199 |  | Plasma Membrane | other |
| GLS | glutaminase | D3Z7P3.1 | -1.115 |  | Cytoplasm | enzyme |
| SLC17A7 | solute carrier family 17 member 7 | BAE34790.1 | 0.579 |  | Plasma Membrane | transporter |
| GRIA2 | glutamate ionotropic receptor AMPA type subunit 2 | AAC37654.1 | 0.905 | Up | Plasma Membrane | ion channel |
| GRIA4 | glutamate ionotropic receptor AMPA type subunit 4 | XP_006509905.1 | 2.008 | Up | Plasma Membrane | ion channel |

#### 3.11.2 ML disease pathway analysis

A major subset of the identified ML disease pathways (Fig 6F) are associated with neurodegenerative and neurodevelopmental disorders, reinforcing the hypothesis that chronic stress accelerates neurodegenerative processes. Top ML disease pathways are presented in fig (Fig 7B). The presence of neuronal ceroid lipofuscinosis, mitochondrial myopathies and encephalopathies suggests that stress-induced molecular changes affect neuronal viability, mitochondrial function, and synaptic integrity. The enrichment of pathways associated with memory deficits, progressive neurological disorders, and motor dysfunction further strengthens the link between chronic stress exposure and cognitive decline. A comprehensive account of extended IPA analysis including detailed canonical pathway analyses, diseases and biofunctions, regulator effects and molecular networks is provided in the Supplementary Information (see Extended Results, Sup Fig. 1-3, and Sup_Table 1-4).

## 4. DISCUSSION

### Sustained Aggression in Parous Females

The control group, consisting of socially housed female mice, exhibited non-aggressive behavior, and was consistent with previous reports on female social dynamics. The absence of aggressive behavior in intact females suggests that mere co-housing with castrated males is insufficient to induce a chronic aggressive state. In contrast to our findings, a study on Swiss Webster (CFW) female mice demonstrated that mere co-housing with castrated males was sufficient to induce aggression towards female intruders (8). Since the lack of aggression in control female CD1 mice was confirmed in our study, we did not include separate control groups for assessments of aggression induction in our modified strategy. Our focus was on developing strategies to enhance aggression in female CD1 mice, more importantly, increase the frequency of aggressive bouts.

Our assessments of postpartum aggression revealed limited and inconsistent aggressive behaviors in female CD1 mice. Although aggression was observed in a subset of individuals, the overall number of aggressive bouts remained low and inconsistent, with a declining trend over consecutive postpartum days. We thus sought an alternative approach by extending the aggression phase through prolonged co-habitation of these CD1 females with the same CD1 males (post castration) that had served as their mating partners prior to parturition. After the co-housing phase, a second aggression assessment was conducted. The comparable cumulative bout counts and similar variance in aggressive behavior by SPF suggest that the modified protocol successfully induces robust and sustained aggression in SPF mice, paralleling the aggression levels observed in male models. Observations on cyclic changes of CD1 females suggest that while estrous cyclicity may influence the propensity for aggression, particularly in the diestrus phase, other factors associated with the experimental paradigm, such as co-housing with castrated males, likely play a significant role in inducing and sustaining aggressive behavior in female CD1 mice.

### Longevity and Reusability of SPF Aggressors Enhance Model Utility

A notable advantage of our model is the demonstrated longevity of the aggressive phenotype in the SPF female aggressors. We observed that the majority of aggressors maintained robust aggressive bouts for at least 120 days (4 months) following the conclusion of a *fem*CSDS paradigm, thereby indicating their potential for serial experimental application. Although formal retesting of aggression was not conducted at 1 or 2-month time point, the sustained aggressive capacity observed at 4 months strongly implies that these SPF females would likely retain, if not fully preserve, their aggressive capabilities at earlier intermediate periods. This enduring stability of the aggressive phenotype substantially augments the utility of our model, facilitating the repeated utilization of a characterized aggressor cohort across multiple *fem*CSDS experiments.

### Behavioral Consequences of *CSDS*: Affective and Anxiety Phenotypes

Defeated mice subjected to *fem*CSDS paradigm exhibited a comprehensive spectrum of depression-like behaviors, including social avoidance, anhedonia, and behavioral despair, consistent with key symptoms of major depressive disorder (MDD) in humans. These findings align with extensive literature from depression models, affirming the face validity of our female model(9–11). Behavioral deficits were accompanied by a significant elevation in serum corticosterone levels, indicative of robust hyperactivation of the hypothalamic-pituitary-adrenal (HPA) axis. Notably, for the current investigation, behavioral and molecular analyses were exclusively conducted on the stress-susceptible population, as determined by the social interaction (SI) ratio.

### Glutamate excitotoxicity signatures

In addition to the NAc, we also assessed glutamate concentrations in the CPu, the dorsal component of the striatum, to evaluate whether the observed excitatory imbalance was localized to the limbic (ventral) striatum or extended across dorsal striatal circuits as well. The CPu is primarily involved in sensorimotor integration, goal-directed behavior, and habit formation, but it also receives inputs from cortical and thalamic regions that are responsive to stress (12,13). We observed significantly elevated glutamate levels in both the CPu and NAc of defeated females. This neurochemical imbalance was paralleled by distinct molecular alterations in the CPu, a robust upregulation of NMDAR2B, a calcium-permeable NMDA receptor subunit, and a concurrent significant reduction in EAAT1, a key astrocytic glutamate transporter. The downregulation of Neurabin, a postsynaptic scaffolding protein, further indicates synaptic instability and structural vulnerability in the CPu.

### Proteomic Insights into NAc Dysregulation in *fem*CSDS induced defeated females

The IPA on NAc proteomics indicated of suppressed Glutamatergic Receptor Signaling Pathway. The pronounced downregulation of SLC1A3 and GLS implies impaired astrocytic uptake and glutamate metabolism, while reduced expression of GRM2, GNG7, GNG12, and GNB5 reflects disrupted metabotropic and G-protein-mediated signaling. In contrast, the upregulation of ionotropic AMPA receptor subunits GRIA2 and GRIA4 may represent a compensatory response to diminished presynaptic control or a maladaptive enhancement of excitatory drive. Findings on glutamate levels in NAc during stress aren’t entirely consistent - some show increases (14,15), some decreases. However studies on addiction predominantly indicate of increased NAc extracellular glutamate levels (16,17).

We focused our proteomic analysis on the NAc because of its central role in the brain’s reward circuitry and its well-established involvement in mood regulation, social behavior, motivation, and susceptibility to stress (18–20). The ESRRA/FOXA1/IL15 regulator effect network reinforces metabolic collapse, showing downregulation of mitochondrial biogenesis regulators. These findings are consistent with the growing understanding of mitochondrial dysfunction as a critical contributor to depression pathophysiology, impacting neuronal excitability and synaptic function.

IPA revealed robust activation of neuroinflammatory pathways, including Neutrophil Degranulation and IL-6 Signaling, with upregulated S100A9 and STAT3 indicating microglial activation and pro-inflammatory cytokine release. Concurrently, the NRF2-mediated Oxidative Stress pathway was significantly implicated, with the suppression of NFE2L2 (Nrf2), a master antioxidant regulator. This, coupled with reduced GPX4 and UCHL1, exacerbates reactive oxygen species (ROS) accumulation and proteasomal dysfunction. The NFE2L2 (Nrf2) regulator effect network further highlights this oxidative burden, which synergizes with mitochondrial dysfunction to amplify synaptic loss and contribute to depressive phenotypes.

### Limitations and future prospects

This study provides significant insights into female-specific stress pathophysiology, yet certain limitations warrant consideration. While IPA provides powerful tools for pathway and network prediction, functional validation of key protein alterations and their causal roles in behavioral deficits would strengthen these findings. Furthermore, while we analyzed certain behavioral outcomes based on estrous cycle stage, our proteomic analysis was not stratified by cyclicity, which is an important consideration given the fluctuating hormonal landscape in females. A crucial future direction involves a direct, parallel proteomic comparison between male and female CSDS model. Our novel *fem*CSDS model successfully recapitulates key behavioral and molecular hallmarks of depression, providing an unprecedented platform to explore sex-specific mechanisms. This work pioneers a path toward equitable, social defeat model. Notably, this model opens new avenues for exploring female aggression, an under-studied area, offering insights distinct from testosterone-based male aggression models.

## Supporting information

Part of the data generated is available in supplementary materials

## Statements & Declarations

## CRediT author statement

**Shashikant Patel:** Conceptualization, Methodology, Investigation, Formal analysis, Data Curation, Writing - Original Draft, Visualization, Writing - Review & Editing ; **Roli Kushwaha:** Investigation, Formal analysis, Data Curation, Visualization, Writing - Original Draft ; **Anusha P.V:** Investigation, Formal analysis, Data Curation ; **Anushka Arvind:** Investigation, Formal analysis, Data Curation ; **Sainath Sunil Dhaygude**: Investigation, Formal analysis, Data Curation; **Satya Ranjan PattnaiK:** Investigation, Formal analysis, Data Curation ; **Arvind Kumar:** Supervision, Resources, Project administration, Writing - Review & Editing ; **Mohammad Idris:** Supervision, Resources, Project administration, Funding acquisition, Writing - Review & Editing; **Sumana Chakravarty:** Conceptualization, Methodology, Investigation, Formal analysis, Data Curation, Supervision, Resources, Project administration, Funding acquisition, Writing - Review & Editing

## Acknowledgements

Authors would like to acknowledge B. Jyothilakshmi and N.Sairam from the Centre for Cellular and Molecular Biology (CCMB), Hyderabad, for the maintenance and care of animals throughout the study period. KIM Department of CSIR-IICT is greatly acknowledged for generating institutional publication number IICT/Pubs./2025/219. The authors acknowledge the use of BioRender.com for the creation of scientific illustrations included in this manuscript.

## Data Availability

Experimental data will be made available on request to corresponding author. Part of the data generated is available in supplementary materials.

