## Supplementary for "Replicating the Gold Standard: A Novel Female Chronic Social Defeat Stress Model (*fem*CSDS) for Studying Sex Differences in Depression"

### 1. Supplementary Methods

#### 1.1 Animals

During the experiments, animals were maintained on a standardized light/dark cycle of 12/12h, with lights on between 06:00-18:00 h and room temperature maintained around 24±1$^{o}$C. Animals were provided food and water ad libitum.

#### 1.2 Experimental strategy

Workflow 1:

We procured intact male and female CD1 mice aged 3-4 months and housed them in groups of four per cage for 7-day habituation (n=12 pairs). Post habituation period, female CD1 mice were housed individually for a week. Concurrently, the male CD1 mice underwent bilateral orchiectomy (castration). Following surgery, the males were housed individually for a week to recuperate, during which the female mice remained individually housed. This ensured that any residual sperm was completely eliminated from the male reproductive system. Post recuperation period, each castrated male mouse was paired with a single intact female CD1 mouse. The pairs were co-housed for 6 weeks to potentially induce aggression in female mice. Aggression assessment was performed for three alternate days consecutively (referred to as Day 1, Day 3 and Day 5) after the co-housing period. To evaluate aggression, the male partner was temporarily removed from the cage for 10 minutes, and a female C57BL/6NCrl mouse was introduced into the female CD1 cage. Aggression was quantified based on behavioral parameters, including aggressive bouts, latency of aggressive bout and offensive sniffing over a 6 minute period. Post aggression assessment sessions the CD1 male partners were returned back to their paired cage. For comparison and to assess aggression induction we used age matched normal CD1 females that didn't underwent any procedure to serve as controls. This protocol (Figure 1: Experimental workflow 1) allowed us to assess whether co-housing with castrated males could induce aggressive behaviors in intact female CD1 mice. In light of our initial findings, we modified our approach to leverage PPA.

#### 1.3 Behavioral tests

##### 1.3.1 Social Interaction Test (SIT)

SIT was conducted on the day after the final defeat session (PDD1). In the first phase, a circular wire-mesh enclosure was positioned at one end of the test arena (60cm x 40cm) without the presence of a CD1 female aggressor. The test mouse was allowed to freely explore the arena for 5 minutes and then was returned to its home cage for a 2-minute interval. In the second phase, the test mouse was reintroduced to the arena for 5 minutes, during which the enclosure contained a novel female CD1 mouse that had not been previously used as an aggressor during the social defeat experiments. The time spent in the interaction zone and the frequency of entries into this zone was evaluated. Mice were segregated into defeated (susceptible) and resilient groups based on SI ratio. The interaction ratio was calculated using the formula:

$$\mathrm{IR} = 100 \times \frac{Time\ spent\ in\ Interaction\ Zone\ with\ Aggressor}{(Time\ \ in\ Interaction\ Zone\ with\ Aggressor + Time\ \ in\ Interaction\ Zone\ without\ Aggressor)}$$

#### 1.3.2 Sucrose Preference Test (SPT)

One week prior to the onset of the stress paradigm, the C57 mice were habituated to a two-choice pipette setup in their home cages, with access to water and 2% sucrose solution. The pipettes were modified to include a drip-free nozzle to prevent spillage. To prevent side bias, the positions of the water and sucrose pipettes were alternated daily throughout the habituation phase. The two choice setup with the pipettes was maintained for the entire 10-day social defeat paradigm. Fluid consumption was recorded on specific days; days 1, 2, 5, 6, 9, and 10. Sucrose preference was calculated using the following formula:

$$Sucrose\ Preference = 100 \times \frac{Volume\ of\ sucrose\ consumed}{Volume\ of\ sucrose\ consumed + Volume\ of\ water\ consumed}$$

#### 1.3.3 Open Field Test (OFT)

To assess anxiety-like behaviors we employed the Open Field Test (OFT), a widely used method for evaluating anxiety and exploratory behavior in rodents. The OFT was conducted in a rectangular arena measuring 40 × 40 × 30 cm, which was virtually divided into central and peripheral zones. Mice were allowed to explore freely in the open field for duration of 5 minutes. During this period, the time spent by each mouse in the central and peripheral zones was recorded. Anxiety-like behaviors were quantified by calculating the frequency of entries into the central zone, the time spent in both the central and peripheral zones and the average velocity of the mice during the exploration period.

#### 1.3.4 Elevated Plus Maze Test (EPM)

The EPM was utilized to evaluate anxiety-like behavior. The EPM consisted of two open arms and two closed arms, allowing for the assessment of exploratory behavior while measuring anxiety levels based on the preference for closed versus open arm. Each mouse was placed in the maze and allowed to explore for 5 minutes. The times spent in the arms were recorded.

#### 1.3.5 Forced Swimming Test

The Forced Swimming Test (FST) was conducted to assess behavioral despair. Mice were individually placed in a transparent, cylindrical glass tank (height: 20 cm, diameter: 15 cm) filled with water maintained around 25°C. Each mouse was allowed to swim for a total of 5 minutes. The latency to immobility and total immobility duration were recorded. Mice were deemed immobile when they adopted a posture characterized by floating motionless in the water, making only the necessary movements to keep their heads above the surface.

#### 1.3.6 Sucrose Splash Test (SST)

The sucrose splash test was used to evaluate self-care and motivational deficits associated with depression-like behavior. A 20% (w/v) sucrose solution was then sprayed onto the dorsal coat using a spray bottle ensuring even coverage without causing stress. The sticky sucrose induces grooming, reflecting motivational drive. Behavior was recorded for 5 minutes using a topmounted camera. Latency to initiate grooming (from spray to first grooming event) and total grooming duration were measured. Grooming was defined as any licking or scratching directed towards the fur or body. Video recordings were scored manually by an independent observer who was blinded to the experimental group allocation.

#### 1.3.7 Vaginal cytology

Smears were obtained using a sterilized pipette filled with 1x PBS, gently flushed against the vaginal wall to collect cells. The collected samples were placed on glass slides, air-dried, and examined under a light microscope for cytological evaluation. The estrous stage was determined based on the proportions of three primary cell types: nucleated epithelial cells, keratinized squamous cells, and leucocytes. Proestrus was identified by the presence of nucleated epithelial cells, while estrus was marked by the dominance of large, keratinized squamous cells. Metestrus (also referred to as early diestrus) was characterized by a mix of leucocytes and a few epithelial cells, whereas diestrus (late diestrus) was primarily defined by the presence of abundant leucocytes. The gradual transitions between these stages were carefully observed, and each mouse was assigned to the appropriate phase for further analysis.

#### 1.3.8 Serum Corticosterone measurement

Corticosterone levels in serum samples were quantified using a competitive ELISA kit (ADI-901-097, Enzo Life Sciences), following the manufacturer’s instructions. Briefly, 10 µL of each serum sample was treated with Steroid Displacement Reagent (1:100 dilution) to dissociate corticosterone from binding proteins. The mixture was incubated, diluted with assay buffer, and added to microplate wells coated with donkey anti-sheep IgG. After incubation with corticosterone-conjugated alkaline phosphatase and corticosterone antibody, plates were washed and incubated with p-nitrophenyl phosphate substrate. The reaction was stopped, and absorbance was read at 405 nm. Corticosterone concentrations were calculated using standard curve.

#### 1.3.9 Glutamate assay

Glutamate levels were measured in serum, NAc and CPu tissue samples using the Glutamate Assay Kit (MAK004, Sigma-Aldrich), following the manufacturer's instructions. Briefly, tissues were homogenized in Glutamate Assay Buffer, and samples were deproteinized to eliminate interfering proteins. Serum samples were directly assayed. Samples and standards were added to a 96-well plate in duplicates, and a colorimetric reaction was initiated by adding Glutamate Developer and Enzyme Mix. After a 30-minute incubation at 37 °C, absorbance was measured at 450 nm using a plate reader. Glutamate concentrations were calculated from a standard curve.

#### 1.3.10 Immunoblotting

Tissue samples were homogenized in 8 M urea lysis buffer supplemented with protease and phosphatase inhibitors. Lysates were sonicated to ensure complete disruption, followed by centrifugation at high speed to remove debris. The resulting supernatants containing total protein were collected, and protein concentrations were quantified using the Bradford assay. Equal amounts of protein (30 µg per sample) were denatured, resolved on SDS-PAGE gels, and transferred onto PVDF membranes. Membranes were blocked and incubated overnight at 4 °C with primary antibodies targeting NMDAR2B, Neurabin, and EAAT1. GAPDH was used as the internal loading control (refer Sup_Table 5 for details). Following primary incubation, appropriate HRP-conjugated secondary antibodies were applied. Densitometric quantification of band intensities was performed using ImageJ software.

#### 1.3.11 Preparation of tissue lysate and Label free Quantitative MS-MS analysis

Samples included 2 control groups and two defeated groups with each group consisting of pooled samples from 4 individual mice. In all samples from 8 control and 8 defeated mice were utilized. Total protein from NAC tissues were extracted using Urea based protein solubilization buffer (1) and quantified using Amido black assay (2) with bovine serum albumin as standard.  Label free Quantitative MS-MS analysis was carried out for all the samples against control. 100 µg of total protein from each experimental group with samples and control tissue lysed protein samples as both experimental and biological duplicates underwent electrophoresis on 10% SDS-PAGE gels, which were then stained with Commassie R250, destained, and the gel was excised into four fractions based on molecular weight. In-gel digestion using trypsin was followed by purification of the digested peptides using C-18 spin columns (Thermo Scientific). The purified peptides were reconstituted in 5% acetonitrile (ACN) and 0.2% formic acid before undergoing Liquid Chromatography/Mass Spectrometry (LCMS/MSMS) analysis using an OrbitrapVelos Nano analyzer (Q-Exactive HF) (3). Proteomic data obtained from the mass spectrometer were analyzed against *Mus musculus* proteome database using Proteome Discoverer 2.2.3 software. The resulting raw data were analyzed using with 1% FDR percolator and XCorr (Score Vs Charge). Differential expressions in proteins were estimated relative to the non-treated control negative samples. Proteins having more than 0.5 log change were recruited for the study.

#### 1.3.12 Brain Area Sampling

Mice were sacrificed by cervical dislocation after behavioral assessments and different brain regions including were microdissected from the freshly collected brain. In brief, the brain was uniformly sliced (1.0mm thickness) on a chilled brain matrix (Harvard Apparatus, USA) after rinsing with ice-cold PBS. NAc was punched with a puncher [12 G] from appropriate slices (4 punches per brain) and snap frozen in liquid nitrogen and stored at -80°C until use.

### 2. Extended Results

#### 2.1 Canonical Pathway Analysis: Synaptic, hormonal and Neurodegenerative Dysregulation in *fem*CSDS.

The Synaptogenesis Signaling Pathway (Sup_Fig 1), exhibited pronounced dysregulation of presynaptic machinery (Sup_Table1). Downregulation of SNAP25 (synaptosome-associated protein 25, Expr Log Ratio= -1.709) and SYN1/2 (synapsin I/II, Expr Log Ratio= -0.849/-0.897) was observed. Compensatory upregulation of STX1B (syntaxin-1B, Expr Log Ratio= 2.8) and VPS18 (vesicular trafficking protein, Expr Log Ratio= 2.7) was also identified. The SNARE Signaling Pathway (p = 8.94E-14), essential for neurotransmitter release, further showed dysregulation of SNAP25 (Expr Log Ratio= -1.709). Oxytocin (OXT) signaling has long been implicated in the regulation of social behaviors, stress responsiveness, and emotional regulation. IPA identified significant dysregulation in the oxytocin signaling pathway, highlighting its potential role in female-specific vulnerability to stress-induced depression. Several G-protein subunits essential for oxytocin receptor coupling, including GNG7, GNB5, GNAI2, and GNA11 (Expr Log Ratios ranging –1.4 to –0.6), were downregulated, suggesting disrupted GPCR-mediated signaling.

Estrogen Receptor Signaling displayed downregulation (ESR1, Expr Log Ratio= 2.1) (Sup_Fig 2). Key proteins in the estrogen receptor signaling pathway were notably altered. mTOR was strongly downregulated (Expr Log Ratio= –2.785), indicating impaired neuroplasticity. Several G-protein subunits (GNG7, GNG12, and GNB5) and kinases like PRKCE also showed reduced levels, suggesting disrupted membrane-initiated estrogen signaling. Conversely, AKT1, AKT3, and PLCL2 were upregulated (Expr Log Ratio= 2.937), pointing to compensatory activation of PI3K-AKT signaling.

#### 2.2 Diseases and Biofunctions Analysis in the Female Chronic Social Defeat Stress Model

The proteomic profile was strongly associated with Neurological Disease ($p$ = 3.42E-56) and Organismal Injury ($p$ = 3.42E-56), reflecting widespread neuronal and systemic stress. Key molecular functions included Cellular Assembly and Organization ($p$ = 2.92E-43) and Nervous System Development ($p$ = 1.01E-42). The enrichment of Mitochondrial Dysfunction ($p$ = 1.84E-12) and NRF2-mediated Oxidative Stress ($p$ = 2.12E-04) pathways were suggestive of metabolic dysregulation in the NAc of defeated mice. Dysregulation of molecules such as CAMK4, CDH2, and GRIA2, which are involved in synaptic remodeling, suggests that depression-related changes in the NAc might extend beyond reward dysfunction and impact broader cognitive and emotional networks. IPA predicted activation state of enriched disease and biofunctions is presented in Sup_Table 2.

#### 2.3 Regulator effects

The ARNT/IPMK/MRTFA network, marked by the highest consistency score (10), highlights disruptions in cytoskeletal dynamics and energy metabolism (Fig 10F). Key targets such as ACTC1, ACTG1 (actin isoforms), and ALDOA (glycolytic enzyme) suggest impaired structural integrity and ATP production, which may contribute to synaptic instability and reduced exploratory behavior. ARNT, a hypoxia-inducible factor partner, and IPMK, a modulator of inositol phosphate metabolism, collectively point to adaptive responses to metabolic stress, potentially compensating for mitochondrial dysfunction. However, sustained suppression of these regulators likely exacerbates neuronal atrophy.

The ESRRA/FOXA1/IL15 network (consistency score = 5.692) (Sup_Fig 3A) further indicates metabolic collapse, with ESRRA (a mitochondrial biogenesis regulator) and FOXA1 (a transcriptional metabolic activator) downregulated alongside targets like ATP5F1B (ATP synthase subunit) and SIRT3 (mitochondrial sirtuin) (Sup_Fig 3A). The NFE2L2 (Nrf2) network (consistency score = -31.197) highlights oxidative stress as a central mechanism. Suppression of NFE2L2, a master antioxidant regulator, coincides with reduced expression of GPX4 (glutathione peroxidase) and UCHL1, exacerbating ROS accumulation and proteasomal dysfunction (Sup_Fig 3E). This oxidative burden may synergize with mitochondrial failure and neuroinflammation, amplifying synaptic loss and depressive phenotypes. Notably, ATG7 upregulation suggests compensatory autophagy, though chronic activation may deplete cellular resources, worsening neuronal atrophy. The top regulator networks are presented in Fig 10 and full list is presented in (Sup_Table 3).

#### 2.4 Network Analysis of Molecular Pathways in the CSDS Model

The IPA of differentially expressed proteins revealed multiple molecular networks with significant enrichment in neurological and developmental functions (Sup_Table 4). Among the analyzed networks, three primary networks designated as Network 1, Network 2, and Network 3 exhibited high relevance to neurobiological processes, with scores of 49, 44, and 42, respectively. Network 1 molecular composition included proteins involved in mitochondrial function, cytoskeletal organization, and metabolic regulation. The prominence of MAPT (tau) suggests a potential vulnerability of microtubule structures to chronic stress. Additionally, the involvement of ribosomal proteins and aminoacyl-tRNA synthetases (ARSs) indicates that stress may disrupt protein synthesis fidelity, affecting synaptic plasticity and neuronal function.

Network 2 demonstrated a functional enrichment in cellular assembly and developmental disorders. The identified molecular interactions suggest a crucial role in maintaining cellular architecture and signaling processes essential for neurogenesis and synaptic maintenance. Key molecules in this network include VCPIP1, SGTA, and BAG6, which regulate protein quality control and degradation via the ER-associated degradation (ERAD) pathway. The involvement of ASPSCR1, UBQLN2, and NPLOC4 suggests alterations in ubiquitin-proteasome system (UPS) activity, a pathway critical for synaptic function and neuronal survival under stress conditions. A key observation in this network is the presence of GABA-B receptor and Kir3 channel, implicating changes in GABAergic signaling.

### TABLES

### Sup_Table 1: Key molecules in the Synaptogenesis Signaling Pathway with their expression log ratios identified through IPA analysis

| Symbol | Entrez Gene Name | GenPept/RefSeq | Expr Log Ratio | Expected | Location | Type(s) |
|---|---|---|---|---|---|---|
| CACNB1 | calcium voltage-gated channel auxiliary subunit beta 1 | BAC80138.1 | -3.658 | | Plasma Membrane | ion channel |
| TLN1 | talin 1 | XP_006537831.3 | -3.462 | Up | Plasma Membrane | other |
| MTOR | mechanistic target of rapamycin kinase | EDL14819.1 | -2.785 | Up | Nucleus | kinase |
| SGTA | small glutamine rich tetratricopeptide repeat co-chaperone alpha | BAC37566.1 | -1.987 | | Cytoplasm | other |
| GRM2 | glutamate metabotropic receptor 2 | EDL21160.1 | -1.888 | Up | Plasma Membrane | G-protein coupled receptor |
| SYT1 | synaptotagmin 1 | NP_001239270.1 | -1.793 | Up | Cytoplasm | transporter |
| CADM1 | cell adhesion molecule 1 | BAD30020.1 | -1.793 | Up | Plasma Membrane | other |
| SNAP25 | synaptosome associated protein 25 | NP_035558.1 | -1.709 | Up | Plasma Membrane | transporter |
| MAPT | microtubule associated protein tau | XP_036012271.1 | -1.447 | Up | Plasma Membrane | other |
| NLGN2 | neuroligin 2 | BAD32437.1 | -1.446 | Up | Plasma Membrane | enzyme |
| PRKCE | protein kinase C epsilon | EDL38612.1 | -1.308 | Up | Cytoplasm | kinase |
| CPLX1 | complexin 1 | EDL20107.1 | -0.995 | Up | Plasma Membrane | transporter |
| SYN2 | synapsin II | NP_001104485.1 | -0.897 | Up | Plasma Membrane | other |
| RAP1B | RAP1B, member of RAS oncogene family | Q99JI6.2 | -0.882 | Up | Cytoplasm | enzyme |
| SYN1 | synapsin I | EDL00736.1 | -0.849 | Up | Plasma Membrane | other |
| CNTNAP1 | contactin associated protein 1 | EDL03896.1 | -0.807 | Up | Plasma Membrane | other |
| ARPC3 | actin related protein 23 complex subunit 3 | BAB22813.1 | -0.794 | Up | Cytoplasm | other |
| STX16 | syntaxin 16 | BAC37129.1 | -0.659 | Up | Cytoplasm | transporter |
| PRKACB | protein kinase cAMP-activated catalytic subunit beta | BAE29331.1 | -0.648 | Up | Cytoplasm | kinase |
| CNTNAP2 | contactin associated protein 2 | BAC98042.1 | -0.646 | | Plasma Membrane | other |
| PAK1 | p21 (RAC1) activated kinase 1 | XP_011239996.1 | -0.629 | Up | Cytoplasm | kinase |
| RRAS2 | RAS related 2 | BAB27607.1 | -0.594 | Up | Plasma Membrane | enzyme |
| PRKAR1B | protein kinase cAMP-dependent type I regulatory subunit beta | AAH11424.1 | -0.589 | Up | Cytoplasm | kinase |
| RAP2A | RAP2A, member of RAS oncogene family | Q80ZJ1.2 | -0.584 | Up | Plasma Membrane | enzyme |
| RAB3A | RAB3A, member RAS oncogene family | P63011.1 | -0.547 | Up | Cytoplasm | enzyme |
| CAMK2D | calciumcalmodulin dependent protein kinase II delta | BAD90304.1 | -0.534 | Up | Cytoplasm | kinase |
| KRAS | KRAS proto-oncogene, GTPase | BAE33023.1 | -0.511 | Up | Cytoplasm | enzyme |
| YKT6 | YKT6 v-SNARE homolog | BAE34934.1 | -0.508 | Up | Cytoplasm | enzyme |
| GUCY1 | guanylate cyclase 1 soluble subunit beta 1 | NP_059497.1 | 0.567 | | Cytoplasm | enzyme |

| | | | | | | |
|---|---|---|---|---|---|---|
| B1 | | | | | | |
| ITSN1 | intersectin 1 | XP_006522994.1 | 0.576 | Up | Cytoplasm | other |
| AP2M1 | adaptor related protein complex 2 subunit mu 1 | BAD32167.1 | 0.599 | Up | Cytoplasm | other |
| NRXN1 | neurexin 1 | BAC41433.2 | 0.643 | Up | Plasma Membrane | transmembrane receptor |
| ITPR1 | inositol 1,4,5-trisphosphate receptor type 1 | XP_006505686.1 | 0.644 | Up | Cytoplasm | ion channel |
| TIAM1 | TIAM Rac1 associated GEF 1 | XP_006523045.1 | 0.644 | Up | Cytoplasm | other |
| SYT5 | synaptotagmin 5 | EDL31245.1 | 0.684 | Up | Cytoplasm | transporter |
| DNAJC5 | DnaJ heat shock protein family (Hsp40) member C5 | EDL07424.1 | 0.699 | | Plasma Membrane | other |
| PIK3R4 | phosphoinositide-3-kinase regulatory subunit 4 | XP_036011263.1 | 0.737 | Up | Cytoplasm | kinase |
| EPHA4 | EPH receptor A4 | CAA46268.1 | 0.785 | Up | Plasma Membrane | kinase |
| BRAF | B-Raf proto-oncogene, serinethreonine kinase | XP_011239436.1 | 0.871 | Up | Cytoplasm | kinase |
| LRP1 | LDL receptor related protein 1 | XP_036011543.1 | 0.894 | Up | Plasma Membrane | transmembrane receptor |
| GRIA2 | glutamate ionotropic receptor AMPA type subunit 2 | AAC37654.1 | 0.905 | Up | Plasma Membrane | ion channel |
| AFDN | afadin, adherens junction formation factor | XP_006523815.1 | 0.907 | Up | Nucleus | other |
| ARHGEF7 | Rho guanine nucleotide exchange factor 7 | XP_036010059.1 | 0.965 | Up | Cytoplasm | other |
| CDH2 | cadherin 2 | AAA37353.1 | 1.039 | Up | Plasma Membrane | other |
| AKT1 | AKT serinethreonine kinase 1 | EDL18587.1 | 1.058 | Up | Cytoplasm | kinase |
| CACNA2D2 | calcium voltage-gated channel auxiliary subunit alpha2delta 2 | AAR89454.1 | 1.109 | | Plasma Membrane | ion channel |
| ADCY5 | adenylate cyclase 5 | AAH90846.1 | 1.22 | Up | Plasma Membrane | enzyme |
| NSF | N-ethylmaleimide sensitive factor, vesicle fusing ATPase | BAC39361.1 | 1.229 | Down | Cytoplasm | transporter |
| AKT3 | AKT serinethreonine kinase 3 | AAH66861.1 | 1.254 | Up | Cytoplasm | kinase |
| GOSR2 | golgi SNAP receptor complex member 2 | AAB82653.1 | 1.402 | Up | Cytoplasm | transporter |
| STXBP2 | syntaxin binding protein 2 | AAH03477.1 | 1.515 | Down | Plasma Membrane | transporter |
| GRIA4 | glutamate ionotropic receptor AMPA type subunit 4 | XP_006509905.1 | 2.008 | Up | Plasma Membrane | ion channel |
| STX1B | syntaxin 1B | BAA25986.1 | 2.838 | Up | Plasma Membrane | other |

**Sup_Table 2: Significantly enriched Diseases and Biofunctions identified in the femCSDS model, reflecting neurological, cellular, and metabolic dysregulation.**

| Categories | Diseases or Functions Annotation | p-value | Predicted Activation State | Activation z-score | Molecules |
|---|---|---|---|---|---|
| Cell-To-Cell Signaling and Interaction,Nervous System Development and Function | Long-term potentiation of hippocampal cells | 0.0000865 | Decreased | -2.433 | CAMK4,CDH2,FMR1,GRIA2,HNRNPK,KRAS,PTPN5,Ptprd |
| Cellular Assembly and Organization,Cellular Compromise | Depolymerization of actin filaments | 0.0000675 | Decreased | -2.183 | CAP1,CFL2,DSTN,GSN,INF2,PFN1,WDR1 |
| Cell Morphology,Connective Tissue Development and Function | Shape change of fibroblast cell lines | 0.0000674 | Decreased | -2.556 | ASAP1,BAIAP2,DGKZ,DOCK3,FLNA,GNA11,GNG12,RAB4A,RHOG,SORBS1,TNIK,TSC2,VIM |
| Energy Production,Molecular Transport,Nucleic Acid Metabolism,Small Molecule Biochemistry | Concentration of ATP | 0.0000426 | Decreased | -2.131 | ACLY,AIFM1,AKT1,ALDOA,ATG2B,ATP5F1B,ATP5IF1,DNM1L,EIF6,GAPDH,GAPDHS,HSD17B10,LDHA,LMNA,LRPPRC,MAOA,MAPT,NAMPT,PLCB3,SIRT2,SIRT3,SLC25A12,SMPD3,YAP1 |
| Developmental Disorder,Neurological Disease,Organismal Injury and Abnormalities,Psychological Disorders | Behavioral deficit | 0.00000326 | Decreased | -2.309 | ABAT,AKT1,ATG7,BRAF,CA2,CDH2,CNTNAP2,CUL3,EIF4E,FMR1,FXR2,GNAI2,GRIA2,IQSEC1,L2HGDH,LRP1,LRRC7,MAOA,MAPT,MECP2,NAMPT,PCDH10,PREP,SHANK2,SHANK3,SIPA1L1,SLC6A1,SLC6A3,TSC2,UBA6 |
| Nucleic Acid Metabolism,Small Molecule Biochemistry | Biosynthesis of ribonucleotide | 0.000000982 | Decreased | -2.041 | ADSS2,AK3,AKT1,ALDOA,AMPD2,ATP5F1B,CAD,CUL4B,DNM1L,DPYSL4,GJA1,HPRT1,KRAS,MAPT,MFN2,NDUFA4,OPA1,PGK1,PRPS2,RPTOR,RYR1,SDHA,SLC25A5,SLC4A7,UBE2O,YWHAZ |
| Cellular Function and Maintenance | Biosynthesis of macromolecule | 0.000000283 | Decreased | -2.175 | ACO1,AKT1,AMPD2,APLP1,CDKN1B,CSDE1,CYFIP1,EEF1A1,EEF1A2,EIF2B1,EIF2S1,EIF3C,EIF3H,EIF3I,EIF3L,EIF3M,EIF4A3,EIF4E,EIF4G1,EIF4G2,EIF4G3,EIF4H,EIF5,EIF5A,EIF6,EPHA4,EPRS1,FMR1,Fus,GAPDH,GFM1,GRM2,GSN,HNRNPD,HNRNPK,HSPB1,ILF3,KHDRBS1,KRAS,MAG,MAPT,MRPL12,MRPL58,MTOR,NPM1,PTK2B,RPL30,RPS20,RPS23,SIRPA,SLC25A5,SMPD3,SNX3,STIP1,SYNCRIP,THY1,TNFAIP8,TP53BP1,TSC2,TXN,UCHL1,UCHL5,UPF1,VIM |
| Behavior | Anxiety | 1.96E-08 | Decreased | -2.545 | AKAP5,ATP1A2,BCAS1,CAMK4,CHL1,COMT,CPNE5,CUL3,DPYSL2,DRG2,EIF4E,FMR1,GNA11,GRIA4,GRM2,ICAM5,LRRC7,MAPT,MECP2,NAMPT,PAK1,PCDH10,PCSK1N,PDYN,Ptma (includes others),PTPRN2,SHANK2,SHANK3,SIPA1L1,SLC17A7,SLC4A10,SLC6A1,SLC6A3,SLITRK1,SV2C |
| Cell-To-Cell Signaling and Interaction,Nervous System Development and Function | Long-term potentiation of hippocampus | 5.71E-09 | Decreased | -2.006 | CAMK4,CAPNS1,CDH2,EPHA4,FMR1,GRIA2,HNRNPK,HSD17B10,ITPR1,ITSN1,KRAS,LRP1,MAPT,PTK2B,PTPN5,Ptprd,SRPK2,STIP1,SYNJ1,SYNPO,TNR,UBQLN2,VPS26A |
| Neurological Disease,Organismal Injury and | Progressive neurological disorder | 3.85E-26 | Decreased | -2.157 | ABAT,ABCD3,ACLY,ACO1,ACTC1,ACTG1,ADAM10,AIFM1,AIMP2,AKAP5,AKT1,ALCAM,ANK3,APLP2,ARHGDIB,ARL6IP5,ASAH1,ASPA,AT |

| | | | | | |
|---|---|---|---|---|---|
| Abnormalities | | | | | AD3A,ATG7,ATL2,ATP1A2,ATP1B1,ATP4A,ATP5PB,ATP6V1B1,ATP6V1E1,BCAS1,BIN1,BRAF,BSG,CELF2,CHL1,CLU,CNKSR2,COMT,COQ7,CYLD,DBNL,DIRAS2,DLAT,DLG2,DLST,DNAJC5,DNAJC6,DNM1L,DPYSL2,Dst,DYNC1H1,EEF1A1,EIF2S1,EIF4G1,ENO2,EPHA4,EPHX1,EPRS1,FH,FIS1,FUBP1,GAD1,GAK,GAPDH,GAPVD1,GBE1,GFAP,GGPS1,GLS,GNAL,GOSR2,GOT2,GRIA2,GRIA4,GSN,HBA1/HBA2,HMGCL,HPRT1,HPX,HSP90AB1,HSPB1,IDE,IGSF8,ITGAV,KCNB1,KHDRBS1,KIF1A,KLC1,KRAS,LDHA,LDHB,LRP1,MACROD1,MAG,MAOA,MAP2K2,MAPT,MATR3,MBP,MDH1,MDH2,MFN2,MMUT,MOG,MTOR,NAXD,NAXE,NEFL,NRCAM,NRXN1,OPA1,OSBPL10,PAK1,PAK3,PDE10A,PDE2A,PFDN1,PFN1,PLD3,PRDX2,PRDX6,PRKACB,PRKCE,PRRC2A,PTGDS,PTGES3,PTPN5,PTPRN2,PTPRZ1,RAB14,RAB39B,RAB3C,RAB6A,RIMBP2,RNF14,RPL13,RTN1,SCFD1,SDHA,SDHB,SERPINE2,SHANK2,SIRPA,SIRT3,SLC1A3,SLC6A3,SNAP25,SPARCL1,SPR,SRPK2,STIP1,SUCLG1,SV2A,SV2C,SYN1,SYN2,SYNJ1,TAGLN,TCEA1,TFRC,THY1,TMED10,TNIK,TNR,TRIO,TUBA1C,TUBA4A,TUBB,TUBB2A,TUBB4A,TXN,UBQLN2,UCHL1,UPF1,VIM,XPO7,YAP1,YWHAZ |
| Cellular Movement,Nervous System Development and Function | Migration of neurons | 0.0000207 | Increased | 2.294 | ABI2,ADAM10,ARHGAP32,CDH2,CHL1,CSDE1,CUL5,DCPS,EPHA4,FBXO41,FLNA,GAD1,GJA1,ITSN1,L1CAM,MAPT,MARK1,MYH10,PTPRZ1,SDC3,SLC1A3,SST,TIAM1,TNR,TRIO,TYRO3,VPS18,YWHAE |
| Neurological Disease,Organismal Injury and Abnormalities | Environmentally induced seizure | 0.000014 | Increased | 2.449 | ATP6V1B2,CNTNAP2,COQ8A,GNG7,SYN1,SYN2 |
| Cellular Movement | Cell movement of neurons | 0.00000718 | Increased | 2.194 | ABI2,ADAM10,ARHGAP32,CDH2,CHL1,CSDE1,CUL5,DCPS,EPHA4,FBXO41,FLNA,GAD1,GJA1,ITSN1,L1CAM,LRP1,MAPT,MARK1,MYH10,NRCAM,PTPRZ1,SDC3,SLC1A3,SST,TIAM1,TNR,TRIO,TYRO3,VPS18,YWHAE |
| Gene Expression | Initiation of expression of RNA | 0.00000211 | Increased | 2 | EIF2B1,EIF2S1,EIF3C,EIF3H,EIF3I,EIF3L,EIF4E,EIF4G1,EIF4G2,EIF4G3,EIF4H,EIF5,FMR1,FUBP1,HSPB1,MECP2,SSRP1 |
| Cellular Function and Maintenance | Receptor-mediated endocytosis | 4.4E-10 | Increased | 2.313 | AAK1,ACSL1,AP2M1,ATP5F1B,ATP6V0C,ATP6V0D1,ATP6V1B2,ATP6V1E1,CAP1,CLTC,CLU,DNAJC6,DNM1,GAK,HGS,HIP1R,HNRNPK,ITSN1,LRP1,NEDD4L,NME1,NSF,PPT1,SF3B3,SFPQ,SHANK3,SYNJ1,SYT1,TFRC,UBQLN2,Wasl |
| Cellular Function and Maintenance | Organization of cells | 1.5E-12 | Increased | 2.236 | ACTG1,ANK3,ATP2B2,BIN1,BSN,C1QA,CAMK1,CFL2,CHL1,CTTNBP2,FLNA,GJA1,GPR158,L1CAM,MAOA,MAPT,MBP,MECP2,MFN2,NEFL,Nefm,NFASC,NLGN2,PAK3,PCLO,RAB39B,RYR1,SLC6A1,SLITRK1,SPARCL1,SPTBN2,SYN1,TNR,TUBB,TXN,WDR1 |
| Behavior | Associative learning | 0.000121 | | | BRSK1,CNTNAP2,MECP2,PPT1,SHANK2,SHANK3,SLC6A1,SNAP25,TNR |
| Nervous System Development and Function | Myelination | 0.000114 | | 0.058 | AKT1,BCAS1,CDH2,CDKN1B,CNTNAP1,CNTNAP2,CYFIP1,HEXA,HGS,HNRNPK,KCNJ10,L1CAM,LRP1,MAG,MAP2K2,MBP,MTOR,PTPRZ1,RPTOR,SIRT2,TPPP,TYRO3,WASF3,Wasl |

### Sup_Table 3: List of Regulator networks

| Consistency Score | Node Total | Regulator Total | Regulators | Target Total | Target Molecules in Dataset | Diseases & Functions |
|---|---|---|---|---|---|---|
| 10 | 33 | 5 | ARNT,IPMK,MRTFA, MRTFB,SKIC2 | 25 | ACTC1,ACTG1,ALCAM,ALDOA,CAPNS1,CD81,CDKN1B,DDAH1,DNAJB4,EIF6,FLNA,GAPDH,HSP90AB1,ITGAV,LDHA,LMNA,MYH10,MYL9,MYO1C,NEO1,PFN1,SORBS1,TFRC,TPI1,VIM | Cell movement,Concentration of ATP,Death of embryo |
| 5.692 | 16 | 5 | ESRRA,FOXA1,IL15,MYC,SKIC2 | 10 | ACLY,AKT1,ALDOA,ATP5F1B,ATP5IF1,GAPDH,LDHA,SIRT2,SIRT3,YAP1 | Concentration of ATP |
| 3.5 | 7 | 1 | FHIT | 4 | IPO7,RAB7A,RANBP1,VIM | Invasion of cells,Migration of tumor cell lines |
| 3.227 | 73 | 8 | ESRRA,MKNK1,MNT, OMA1,RORC,TFEB,TRIB1,TSC1 | 60 | ACADVL,ACSL1,ACSL6,ACTC1,ALCAM,ALDOA,ALDOC,AP1B1,ASAH1,ATP1B1,ATP5F1B,ATP6V0A1,ATP6V0C,ATP6V0D1,ATP6V1A,CACNA2D2,CACNB1,CAD,CLTA,CLTC,CLU,CRMP1,DNM1L,FLNA,GAA,GAPDH,GLS,GOT1,GOT2,HADHA,ITPR1,KLC1,LDHA,LDHB,MAPT,MDH1,MDH2,MFN2,NDUFA4,NRCAM,NRXN1,OPA1,PGAM1,PGK1,PPT1,RAB14,RAB3A,RAB7A,SDHB,SIRT3,SLC1A3,SNAP25,SPARCL1,SYN2,TAGLN,TPI1,TSC2,UBE2O,VIM,YWHAZ | Biosynthesis of ribonucleotide,Gluconeogenesis,Infection by RNA virus,Macrovesicular hepatic steatosis,Progressive neurological disorder |
| 3.015 | 16 | 3 | CLUH,LARP1,SRC | 11 | CDKN1B,EEF1A1,EIF3H,FH,GFM1,OPA1,RPL30,RPS20,RPS23,SIRPA,VIM | Death of embryo,Synthesis of protein |
| 1.768 | 11 | 2 | NR6A1,UBA1 | 8 | CAMK1,CDKN1B,GLS,HSPB1,LMNA,NEFL,TAGLN,VIM | Apoptosis |
| 1.155 | 5 | 1 | SP1 | 3 | FLNA,PRKCE,VIM | Cell spreading of kidney cell lines |
| 0.603 | 18 | 6 | ADORA2A, CD40,INSR,PALMD,RORC,STK11 | 11 | ALDOC,GAPDH,GOT1,HADHA,MDH1, MDH2,MFN2,PGAM1,PGK1,PGM1,TPI1 | Gluconeogenesis |
| 0.13 | 63 | 1 | MAPT | 59 | AAK1,ACLY,ACTG1,ASAH1,ATP1A2,ATP1B1,ATP4A,ATP5F1B,ATP5PB,ATP6V0D1,ATP6V1E1,BSG,BSN,CAP1,CFL2,CLTC,DNAJC5,DNAJC6,DNM1L,DPYSL2,DYNC1H1,ENO2,EPHX1,EPRS1,GAD1,GAPDH,GFAP,GOT2,HBA1/HBA2,HSPB1,KLC1,LDHB,LRP1,MBP,MDH1,MOG,MTOR,NME1,PAK1,PAK3,PCLO,PDE2A,PPT1,PRDX6,SLC1A3,SNAP25,STIP1,SUCLG1,SYN1,SYNJ1,THY1,TUBA1C,TUBA4A,TUBB,TUBB2A,TXN,UCHL1,VIM,YWHAZ | Organization of cells,Progressive neurological disorder,Receptor-mediated endocytosis |
| -1.923 | 55 | 1 | MAPT | 53 | AAK1,ACAT1,ACLY,ALDOA,ASAH1,ATP1A2,ATP5F1B,BSG,CAMK2D,CDKN1B,CLTC,CYFIP2,DNAJC5,DNM1L,DYNC1H1,EPHX1,EPRS1,Fus,GAPDH,GFAP,GOT2,GPX4,HBA1/HBA2,HK1,HSPB1,HSPH1,INPP4A,LRP1,MBP,MDH1,MOG,MTOR,NME1,PAK1,PAK3,PPP2CA | Necrosis |

| | | | | | | |
|---|---|---|---|---|---|---|
| | | | | | ,PPT1,PRDX3,PRDX6,PTK2B,SLC1A3,SNAP25,STIP1,STX1B,THY1,TLN1,TUBB,TXN,UCHL1,UQCRQ,VIM,YWHAE,YWHAZ | |
| -1.938 | 47 | 1 | MAPT | 45 | ACLY,ACTG1,ADD1,ADD2,ALDOA,ANXA6,ATP5F1B,BSG,CAMK2D,CAP1,CDKN1B,CLTC,DNM1L,DPYSL2,DYNC1H1,EPRS1,FBXO41,GAD1,GAPDH,GFAP,GOT2,GPX4,GUCY1B1,HK1,HSPB1,INPP4A,LRP1,Mapre2,MOG,MTOR,NME1,PAK1,PAK3,PDE2A,PTK2B,SEPTIN7,SLC1A3,SNAP25,THY1,TLN1,Tpm1,TXN,VIM,YWHAE,YWHAZ | Migration of cells |
| -2.846 | 12 | 1 | RORC | 10 | ALDOA,GLS,GLUD1,GOT1,GOT2,LDHA,MDH1,NSDHL,PGK1,UBE2O | Cell proliferation of tumor cell lines |
| -4 | 11 | 1 | TFEB | 9 | AP1B1,ATP6V0A1,ATP6V0C,ATP6V0D1,ATP6V1A,CLTA,CLTC,RAB7A,VIM | Infection of cells |
| -4.491 | 8 | 1 | STX18 | 6 | CADM1,ILF3,SFPQ,SPOCK1,SSRP1,TAGLN | Apoptosis |
| -6 | 6 | 1 | CDKN2B | 4 | GLS,GLUD1,GOT1,GOT2 | Cell proliferation of tumor cell lines |
| -6.708 | 7 | 1 | SP1 | 5 | CDH2,GNAI2,GRIA2,MAOA,MECP2 | Behavioral deficit |
| -10.205 | 9 | 1 | MKNK1 | 7 | CLU,CRMP1,MAPT,NCDN,NRCAM,NRXN1,VIM | Outgrowth of cells |
| -12.333 | 12 | 1 | CD40 | 10 | AIMP2,CDKN1B,CUL1,GAPDH,GLS,HK1,IPO4,LDHA,NPM1,UQCRFS1 | Cell death of tumor cell lines |
| -12.969 | 7 | 1 | CHI3L1 | 5 | AKT3,LDHA,NAMPT,TPPP,VIM | Cell movement of tumor cell lines |
| -13.5 | 18 | 1 | SP1 | 16 | BSG,CDKN1B,CPLX1,DLAT,DNM1,GJA1,GOT1,GRIA2,ITGAV,ITPR1,MAOA,MFN2,PRKCE,RANBP1,SERPINE2,SLC25A5 | Apoptosis of tumor cell lines |
| -31.197 | 31 | 1 | NFE2L2 | 29 | ATG7,CBR1,CCT3,CUL1,DCTN3,EIF2S1,EIF3C,EIF4G2,GBE1,GLS,GLUD1,GNA11,GOT1,GPX4,HSP90AB1,IDE,L1CAM,LDHA,LMNA,MAPT,PCDH7,PSMD3,PSMD7,RANBP1,SIRT3,STIP1,TP53BP1,TXN,UCHL1 | Cell death of tumor cell lines |

**Sup_Table 4: Top Molecular Networks, score and their associated neurological and developmental functions identified through IPA analysis**

| ID | Molecules in Network | Score | Focus Molecules | Top Diseases and Functions |
|---|---|---|---|---|
| 1 | 39S ribosomal subunit,ABHD10,ACSF3,ACTC1,ADD2,AIMP1,AIMP2,CCDC124,CPNE6,EPRS1,FUBP1,glutamine-tRNA ligase,HNRNPUL2,IARS1,ITFG1,LAMP2A trimer,LARS1,LARS2,MAP7D1,MAPT,MAPT-8:Microtubule,MATR3,MRPL12,PPT1,PSPC1,Ptma (includes others),Qars1,RTRAF,SEPTIN6,SEPTIN7,SFXN5,SPOCK1,TPPP,YLPM1,ZFR | 49 | 31 | Developmental Disorder, Neurological Disease, RNA Post-Transcriptional Modification |
| 2 | ACSL3,AMPD2,ASPSCR1,BLMH,CSDE1,DNPH1,FAF2,GABA B receptor G-protein beta-gamma and Kir3 channel,GET3,GET4,HEXA,HSPA8:LAMP2a multimeric,IDI1,KCNJ10,LAMP2a multimer complex:GFAP,LRRC40,Mislocalized membrane protein:SGTA:BAG6:GET4:UBL4A:ASNA1:ATP,MTO1,NARS1,NMRAL1,NPLOC4,NSDHL,PPP1R11,SGTA,SGTB,SLITRK1,SREBP,Tail-anchored protein:SGTA:BAG6:GET4:UBL4A:ASNA1:ATP,TKFC,TRMT112,UBQLN2,UBXN2B,VCPIP1,VIM,WDR48 | 44 | 29 | Cellular Assembly and Organization, Developmental Disorder, Protein Synthesis |
| 3 | ACAC,ACBD3,ADD1,ALCAM,ARFIP2,ARL1,ATG,ATG2B,beta-galactosidase,C2CD2L,CADPS,cis-SNARE:3xSNAP:NSF hexamer,CISD2,GGC-RAB4A:GTP:KIF3:microtubule,INF2,KRAS,LLGL1,MOCS2,MYADM,NEO1,PCDH10,PLLP,RAB35,RAB4A,RAB7A,RABGAP1,RAC2 plasma membrane effectors,RAC2:GTP:RAC2 cytosolic effectors,RIMOC1,SLC1A3,SLC24A2,SLC30A1,SV2A,TPD52L2,YKT6 | 42 | 28 | Embryonic Development, Nervous System Development and Function, Organ Development |
| 4 | 3-hydroxyacyl-CoA dehydrogenase,ACADVL,ACAT1,AIFM1,AK3,CALCOCO1,DHRS7B,DLST,EEF1D,EIF5A,GABARAPL1,GJA1,GLUD1,GOT2,GPX4,growth hormone,HADH,HADHA,HSD17B,HSD17B10,HSD17B12,IL12 (family),LDL/cholesterol,MDH2,mediator,NAMPT,PREP,proinsulin,PTPRN2,SGSM1,SIRT3,SUCLG1,UAP1L1,VPS52,WDR82 | 42 | 28 | Developmental Disorder, Hereditary Disorder, Lipid Metabolism |
| 5 | alpha catenin,ATAC B,ATAC C,ATAD3A,caspase 3/7,DLAT,EFTUD2,EIF6,EPB41L3,FBL,GFAP,HNRNPH1,HNRNPK,IPO7,NME1,PDIA4,PDK,PI3K p85,PRDX6,PRMT5,PRPS2,RAB11FIP5,RNF11,RSK,SF3B3,SMAD2/3,STRAP,SYK/ZAP,SYNCRIP,TFRC,TIAM1,TYRO3,UQCRQ,YAP1,YARS1 | 37 | 26 | [Cellular Development, Cellular Growth and Proliferation, RNA Post-Transcriptional Modification |
| 6 | ACLY,ALDOA,ALDOC,AMPK,ARMC8,ASAH1,ATG7,ATP5IF1,C6orf136,CAB39,CAND1,CK2:FACT,CRL4 E3 ubiquitin ligase:CAND1,CSNK2B,CUTA,DBNL,EEF1A1,Ficolin-rich granule lumen proteins,FLG fragment:Keratin tonofilament:Desmosome,GSN,GYG1,HPRT1,Keratin tonofilament:Desmosome,KPNB1,KRT1,MAP1S,MEF2,OSCP1,PAFAH1B2,PDAP1,PGAM1,PGM1,Secretory granule lumen proteins,Specific granule lumen proteins,Tertiary granule lumen proteins | 35 | 25 | Cancer, Organismal Injury and Abnormalities, Reproductive System Disease |
| 7 | ACOT11,BAIAP2,CAMK2D,CEP170,CIT,CNTNAP5,CYFIP2,DLG2,DMWD,Fbll1,FMR1,Gly,D-Ser:L-Glu:GRIN1:GRIN2 NMDA receptors,Gly,D-Ser:L-Glu:GRIN1:GRIN2 NMDA receptors:CALM1:4xCa2 ,Gly,D-Ser:L-Glu:GRIN1:GRIN2 NMDA receptors:PSD proteins:Mg2 ,Gly,D-Ser:L-Glu:GRIN1:GRIN2B NMDA receptor:RASGRF1 oligomer:CALM1:4xCa2 ,Gly,D-Ser:L-Glu:GRIN1:GRIN2B NMDA receptors,Gly,D-Ser:L-Glu:GRIN1:GRIN2B NMDA receptors:CaMKII dodecamer:CALM1:4xCa2 ,GRIA2,GRIN1:GRIN2 NMDA receptors:PSD proteins,GRIN1:GRIN2 NMDA receptors:PSD proteins:Mg2 ,IQSEC1,KHDRBS1,LRRC7,MAP7,NEFL,NLGN2,P-TEFb,PCLO,PRRC2A,RAS GEFs,SHANK3,SHISA7,SLC4A10,SRPK2,TNIK | 35 | 25 | Nervous System Development and Function, Organismal Development, Tissue Development |

**Sup_Table 5: List of antibodies used in the study**

| Target Protein | Antibody Type | Host Species | Source / Company | Catalog Number | Dilution |
|---|---|---|---|---|---|
| NMDAR2B (GRIN2B) | Primary | Abcam | Ab65783 | Ab65783 | 1:1000 |
| Neurabin | Primary | CST | 4166S | 4166S | 1:1000 |
| EAAT1 | Primary | Abcam | Ab8245 | Ab8245 | 1:1000 |
| GAPDH | Primary | Abcam | Ab18561 | Ab18561 | 1:1000 |
| Anti-Rabbit IgG | Secondary | Invitrogen | G21234 | G21234 | 1:5000 |
| Anti-Mouse IgG | Secondary | Invitrogen | G21040 | G21040 | 1:5000 |

### FIGURES

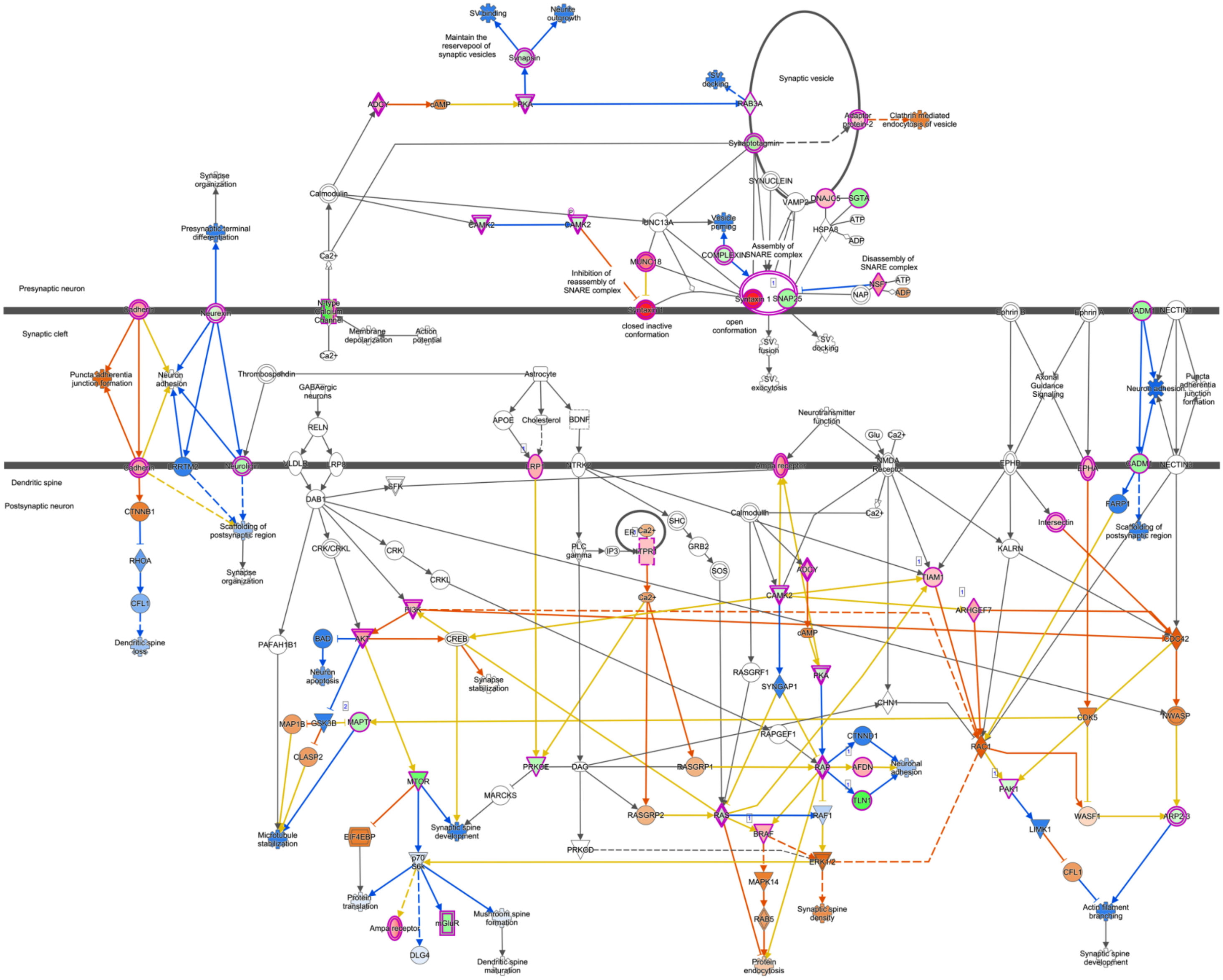

**Sup_Fig 1:** Dysregulation of the Synaptogenesis Signaling Pathway in femCSDS. The graphical representation visualizes expression changes of molecules: red nodes indicate increased expression, and green nodes denote decreased expression, with color intensity reflecting the magnitude of the change. Within this pathway, core presynaptic components crucial for neurotransmitter release were downregulated, including SNAP25 (log ratio = -1.709), SYN1 (log ratio = -0.849), and SYN2 (log ratio = - 0.897), suggesting impaired vesicle fusion. Counteracting this, STX1B (Syntaxin 1B), also involved in synaptic vesicle fusion, showed a substantial upregulation (log ratio = 2.838), potentially indicating a compensatory response or altered synaptic organization. Further, MTOR, a central regulator of synaptic plasticity, was significantly downregulated (log ratio = -2.785) , while AMPA receptor subunits GRIA2 (log ratio = 0.905) and GRIA4 (log ratio = 2.008) were upregulated, suggesting altered postsynaptic sensitivity. Other critical changes included decreased CACNB1 (log ratio = -3.658) and TLN1 (log ratio = - 3.462) impacting neuronal excitability and cytoskeletal integrity. Predicted relationships between molecules are shown by lines, with solid lines indicating direct and dashed lines indirect interactions, and colors denoting predicted activation (orange) or inhibition (blue). These changes highlight a profound dysregulation of synaptic architecture, neurotransmitter release machinery, and plasticity mechanisms in defeated female mice, contributing to the observed behavioral deficits.

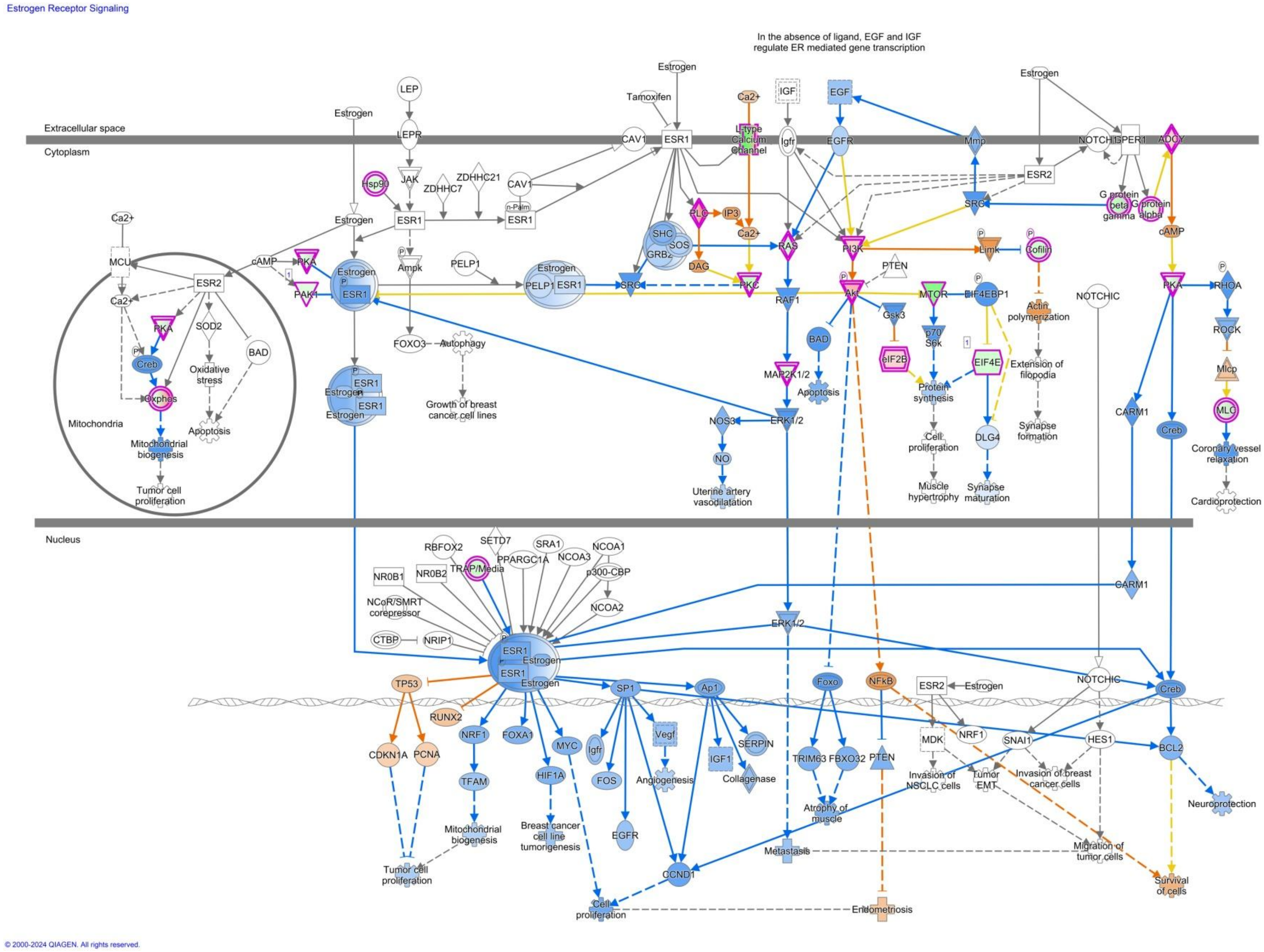

© 2000-2024 QIAGEN. All rights reserved.

**Sup_Fig2: Estrogen receptor signaling pathway**

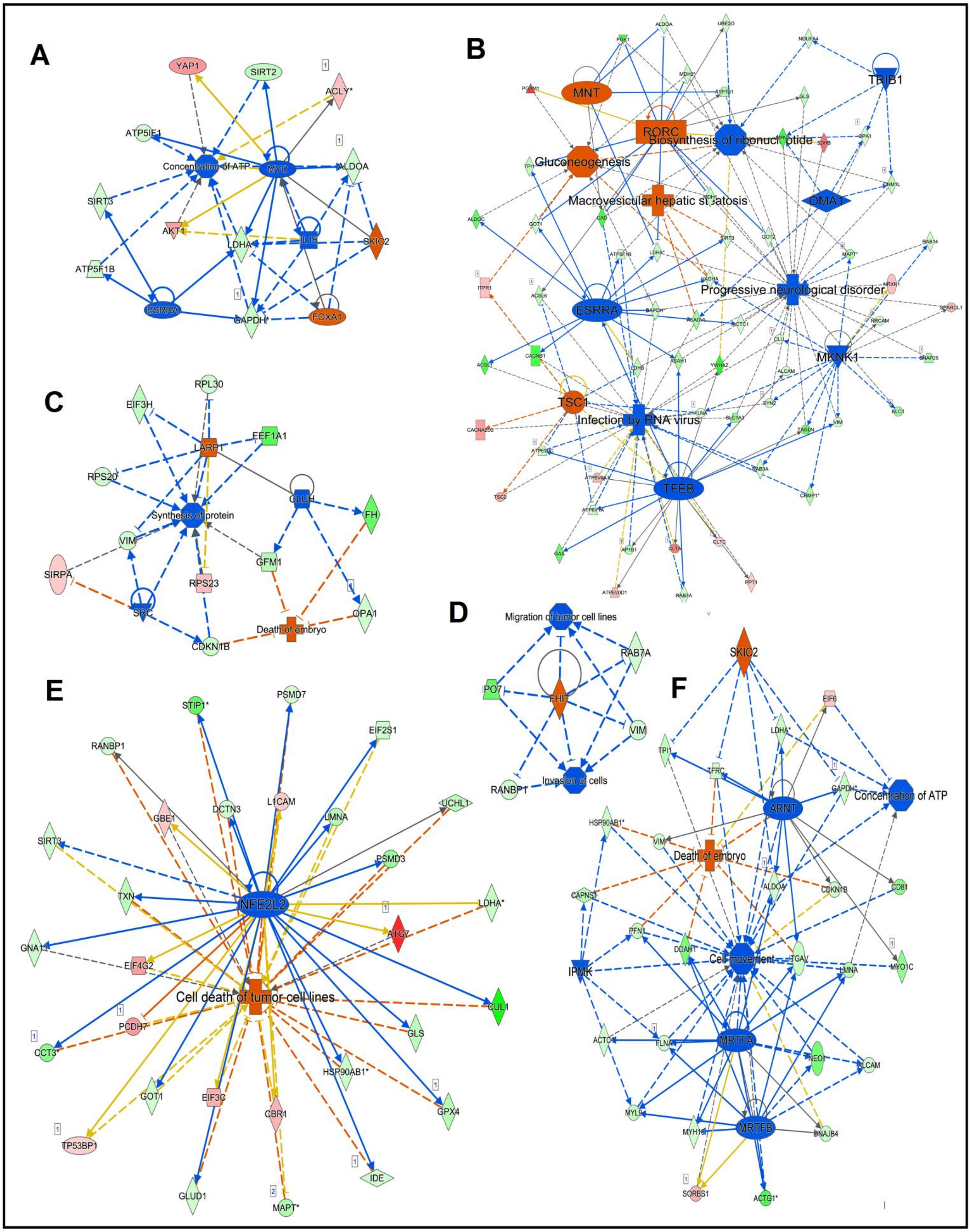

A
YAP1
SIRT2
ACLY*
ATP5IF1
Concentration of ATP
MYC
ALDOA
SIRT3
AKT1
LDHA*
IL15
SKIC2
ATP5F1B
ESRRA
GAPDH*
FOXA1
B
MNT
RORC
TRIB1
Gluconeogenesis
Biosynthesis of ribonucleotide
Macrovesicular hepatic steatosis
OMA1
ESRRA
Progressive neurological disorder
MKNK1
TSC1
Infection by RNA virus
TFEB
C
RPL30
EIF3H
EEF1A1
LARP1
RPS20
CLUH
Synthesis of protein
FH
VIM
GFM1
SIRPA
RPS23
SRC
OPA1
Death of embryo
CDKN1B
D
Migration of tumor cell lines
RAB7A
IPO7
FHIT
VIM
RANBP1
Invasion of cells
E
STIP1*
PSMD7
EIF2S1
RANBP1
L1CAM
DCTN3
UCHL1
GBE1
LMNA
SIRT3
PSMD3
NFE2L2
TXN
LDHA*
ATG7
GNA11
EIF4G2
Cell death of tumor cell lines
CUL1
PCDH7
CCT3*
GLS
HSP90AB1*
GOT1
EIF3C
CBR1
GPX4
TP53BP1
IDE
GLUD1
MAPT*
F
SKIC2
EIF6
LDHA*
TPI1
TFRC
GAPDH*
Concentration of ATP
ARNT
HSP90AB1*
VIM*
Death of embryo
CAPNS1
ALDOA
CDKN1B
CD81
PFN1
DDAH1*
Cell movement
ITGAV
LMNA
MYO1C
IPMK
ACTC1
FLNA
MRTFA
NEO1
ALCAM
MYL9
MYH10
MRTFB
DNAJB4
SORBS1
ACTG1*

**Sup_Fig 3: Ingenuity Pathway Analysis (IPA) Regulator Effects.** (A) ESRRA/FOXA1/IL15 Network (consistency score = 5.692) highlights metabolic collapse through the downregulation of mitochondrial biogenesis regulators and ATP synthase components. (B) ESRRA, MKNK1, MNT, OMA1, RORC, TFEB, TRIB1, TSC1 Network shows distinct metabolic and protein synthesis regulation. (C) CLUH, LARP1, SRC Network details interactions related to protein localization and cell signaling. (D) FHIT Network focuses on its predicted regulatory influence on cellular processes. (E) NFE2L2 (Nrf2) Network (consistency score = -31.197) points to oxidative stress, showing suppression of this master antioxidant regulator and its targets, leading to increased ROS and proteasomal dysfunction. (F) ARNT/IPMK/MRTFA Network (consistency score = 10) indicates disruptions in cytoskeletal dynamics and energy metabolism, affecting structural integrity and ATP production.
